# Mechanisms of antibody binding revealed by asymmetric Fab-virus complexes

**DOI:** 10.1101/2020.12.01.406983

**Authors:** Daniel J. Goetschius, Samantha R. Hartmann, Lindsey J. Organtini, Heather Callaway, Kai Huang, Carol M. Bator, Robert E. Ashley, Alexander M. Makhov, James F. Conway, Colin R. Parrish, Susan Hafenstein

## Abstract

Overlap on the surface of parvovirus capsids between the antigenic epitope and the receptor binding site contributes to species jumping. Mab 14 strongly binds and neutralizes canine, but not feline parvovirus. The high resolution map of the canine parvovirus capsid complexed with Fab 14 was used to solve local structures of the Fab-bound and -unbound antigenic sites extracted from the same complex. The subsequent analysis includes a new method for using cryo EM to investigate complementarity of antibody binding.

## Introduction

Canine parvovirus (CPV) emerged as a variant of a virus in the mid-1970s, subsequently causing a pandemic of disease in dogs during 1978 (1, 2). Since that time, multiple variants have emerged with additional mutations in the viral capsid (3, 4). Extensive genetic and biochemical studies have shown that specific mutations displayed on or near the capsid surface alter the CPV binding phenotype to the host receptor, transferrin receptor type-1 (TfR). Since the specific host ranges of canine and feline parvoviruses is primarily controlled by the ability of the virus to bind TfR, changes in the binding site alter the ability of the virus to infect different hosts (3, 5, 6).

The virus capsid is highly antigenic, and an infection elicits many different host antibodies, which recognize specific structures on the surface of the virus that are primarily displayed as conformational epitopes. Many antibodies efficiently neutralize virus as IgGs, whereas they vary in their neutralization abilities when tested as Fabs (7, 8). In a number of cases selection of antibody escape mutations by antibodies also selects for host range variation in the viruses, while selection for host range variation also alters the antigenic structure recognized by specific antibodies. Although these changes appear to result from overlap of the receptor and antibody binding sites, it is still not clear how different selections may operate in the natural evolution of the viruses. Understanding the mechanisms of host recognition and the dynamics of the binding by antibodies and receptors would allow us to predict the ability of a given virus capsid to change hosts or to escape host immunity, and to make the connections between those processes.

CPV has a small, 26 nm diameter, T=1 icosahedral capsid that packages a single-stranded DNA genome of about 5,000 bases. The capsid shell is comprised of VP1 (90%) and VP2 (10%), which are generated by differential mRNA splicing events so that the entire sequence of VP2 is also contained within VP1. Both proteins fold into the same eight-stranded, anti-parallel β-barrel structure, where the β-strands are connected by loops that make up the surface features of the capsid. A raised region known as the threefold spike surrounds each icosahedral three fold axis and contains most of the antigenic structures recognized by different antibodies (9, 10). MAb 14 is a mouse monoclonal antibody generated against CPV capsids that has particularly interesting properties. MAb 14 binding, hemagglutination inhibition (HAI) properties, and neutralization are all virus-strain specific, and it bound with significantly higher affinity to CPV capsids than to the closely related but host range variant virus that infected cats, feline panleukopenia virus (FPV) (11–13). The virus-specific binding of MAb is controlled by the capsid surface residue 93, which is Lys in FPV and Asn in CPV (14–16). In addition to antibody recognition, residue 93 also controls canine host range, since Asn93 allows binding to canine TfR and infection of canine cells, whereas Lys93 in the equivalent position on the FPV capsid prevents both of these processes (15).

Despite the central role of antibodies in protecting animals against virus infections and allowing recovery from disease, in many cases we still lack a detailed understanding of epitope characteristics, the dynamics of binding processes, and the viral neutralization mechanisms. Previous X-ray crystallography and cryo-electron microscopy (cryo-EM) structures include the Fab of MAb14 (Fab 14), virus capsids, and a Fab 14-CPV capsid complex at moderate resolution (PDB IDs 2CAS, 1C8F and 3GK8) (17–19). Crystal structures of Fab 14, CPV, and FPV fitted into the cryo-EM map of Fab-virus complex have allowed us to predict protein interactions in the binding interface (19). Although this was the most rigorous approach at the time, the resulting pseudo-atomic structure based on the fitting did not explain why variation in residue 93 controlled Fab binding or identify likely mechanisms of antibody neutralization. Recent technological advances in cryo-EM now allow us to solve Fab-virus structures at high enough resolution to build atomic models directly into the density map for identifying interactions unambiguously.

The binding and occupancy of Fabs on parvovirus have also been determined previously using charge detection mass spectrometry (CDMS), which revealed that some of the tested monoclonal antibody-derived Fabs, including Fab 14, could fully occupy all 60 epitopes of the capsid, but with some differences in the kinetics of attachment (20). Incubating with excess Fab molecules to occupy all icosahedrally-equivalent sites on capsids has long been the preferred cryo-EM structural approach since this allows icosahedral symmetry averaging to be imposed during the reconstruction process for maximizing resolution (19, 21, 22). However, there are few studies confirming the occupancies of Fabs on viral capsids, and the IgG likely does not saturate the entire surface of the capsid, so that an undersaturated capsid (with fewer than 60 bound Fab, in the case of parvoviruses) would more closely mimic a physiologically relevant setting. Solving such an asymmetric structure at atomic resolution is now possible due to advances in cryo-EM and the reconstruction approaches.

Here we define an atomic model of Fab 14 bound to the capsid of CPV based on cryo-EM of the complex, and examine the functional mechanisms that affect binding by testing antibody mutants. Of the two data sets used to reconstruct Fab-virus complex maps, one had close to complete occupancy of the 60 capsid epitopes, whereas the other had an average of 10 Fabs bound per capsid. These data were used to solve the icosahedrally averaged structures of fully Fab-occupied and partially Fab-occupied complexes to resolutions of 3.2 Å and 2.3 Å, respectively. An asymmetric, partially Fab-occupied virus map calculated with local reconstruction approaches attained 2.4 Å global resolution and revealed the Fab-occupied and unoccupied sites on the same virus capsid. These structures allowed unambiguous identification of residues and side chains involved in the Fab-virus binding interface and also revealed local conformational changes in the antibody binding site and capsid epitope induced by Fab binding. The partial occupancy of the capsids by Fab also provided an opportunity to develop innovative algorithms to test for complementarity of Fab binding to different positions on the capsid.

Notably, it was the asymmetric approach and not the traditional icosahedrally averaged reconstruction that revealed the mechanism of virus strain-specific attachment and neutralization.

## METHODS

### Production of viruses, antibody, and Fab

Viral capsids were produced as previously described (23, 24). Briefly, Norden Laboratory Feline Kidney (NLFK) cells were infected with FPV or CPV and incubated for 5 days. The culture supernatants were collected and clarified by centrifugation at 10,000 RPM for 15 minutes, then capsids were precipitated overnight with polyethylene glycol 6000, resuspended and banded in a linear 10% to 30% sucrose gradient at 100,000 × g for 3 hours, then full and empty capsids were collected separately. The IgGs were purified from hybridoma supernatants by Protein G chromatography, and the Fab isolated after digestion with pepsin as described previously (8). Briefly, the Fab fragment was removed by protein G binding, and the monomeric Fab isolated by size-exclusion chromatography in an S100 column.

### Sample preparation and data collection

Two preparations of virus capsids and Fab were incubated at room temperature for 1 hour to produce fully occupied Fab and partially occupied Fab complexes, designated full-Fab and low-Fab, respectively. Three μl of each incubation was applied to separate Quantifoil grids (Quantifoil Micro Tools GmbH, Jena, Germany), blotted to remove excess, and plunge-frozen into a liquid ethane-propane mixture or ethane alone using an Mk III Vitrobot (FEI, Hillsboro, OR). For the full-Fab data, the sample was applied to R2/1 grids coated with a 2nm-layer of continuous carbon, and low-dose micrographs were recorded using an FEI Polara G2 microscope operating at 300 kV in nanoprobe and a nominal magnification of 78,000x with defocus values ranging from −0.5 to −4.2 μm. Images were collected under the software control of the EPU program using an FEI Falcon 3 direct electron detector (DED) with post-column magnification of 1.4x yielding a calibrated pixel size at the sample of 1.35Å (**Table S1**). For the low-Fab sample, data were recorded using an FEI Titan Krios microscope operating at 300 kV and a nominal magnification of 59,000x with defocus values ranging from −0.7 to −4.9 μm. Images were also collected under control of EPU on a Falcon 3 DED with a calibrated pixel size at the sample of 1.1Å. Both data sets were recorded in movie mode by recording multiple frames corresponding to one field, allowing for correction of beam-induced movement.

### Reconstruction approaches

#### Icosahedral reconstruction

Particles were autopicked using manually selected templates and contrast transfer functions (CTF) were estimated using GCTF (25). RELION was used for motion correction, movie refinement, and particle polishing (26), whereas cryoSPARC was used for particle sorting and high resolution icosahedral refinement (27). Ctf refinement with correction for higher order aberrations was performed in RELION. Atomic models were built using Coot (28) and Phenix (29), using crystal structures 2CAS (CPV) and 3GK8 (Fab 14) as starting models (17, 19), before validation in MolProbity (30).

#### Localized classification

Subparticle classification was done with localized reconstruction scripts and 3D classification in RELION (26, 31). The subparticle was defined by docking a crystal structure of Fab 14 (PDB ID 3GK8) into the density map (19), and extracted using ISECC (32). These subparticles then underwent 3D classification in RELION without translations or rotations. This allowed for discrimination between Fab-occupied and -unoccupied subparticles. Subparticles were classified into six classes, resulting in strict distinction between occupied and unoccupied epitopes. There was a diversity of unoccupied states, featuring varying amounts of Fab density from the immediately adjacent epitopes (**S.Fig. 1**).

#### Asymmetric reconstructions

After 3D classification into 6 subparticle classes to distinguish between Fab-occupied and -unoccupied subparticles, a symmetry-break operation was accomplished using ISECC_symbreak. For the two states, Euler angles for each subparticle were reassigned to their corresponding whole complex image. This produced two separate data files containing particle orientations (.star files), corresponding to an either Fab-occupied or -unoccupied state at a selected site. Even though there were far more Fab-unoccupied sites, in order to make valid comparisons an equal number of Fab-occupied and Fab-unoccupied subparticles were selected from each particle. This process, which we termed “particle matching”, ensured equivalent scaling of both the occupied and unoccupied maps (**S.Fig. 1**). Both maps contained the same particle images, in identical numbers, differing only in the orientations. Orientations were derived from strict symmetry-expansion of the original icosahedral refinement, without local refinement, ensuring equivalent accuracy angles in the final maps. This matching approach provided equal data sets for a total of 1,657,372 particle-orientations per reconstruction. DeepEMhancer was used to improve local sharpening of maps during post-processing (33). Difference maps were calculated by subtracting the Fab-occupied and unoccupied maps in EMAN (34).

#### Correlated local classification

To describe the configuration of individual bound Fab molecules on each Fab-virus complex, we implemented a system analogous to a radial distribution function (RDF). Each component of the RDF represents the 3D distance between occupied sites on the capsid, as determined by 3D classification and the vectors used in subparticle extraction. On a per-virus basis, the distance between each pair of subparticles was calculated, yielding a list of (n^2^-n)/2 distances, where n = the number of Fab molecules bound to the given complex. Fab binding patterns were derived by normalizing the observed RDFs to a hypothetical particle with all binding site occupied. This combined novel approach was termed correlated local classification. The custom software package, ISECC (icosahedral subparticle extraction & correlated classification), is available for download at https://github.com/goetschius/isecc.

#### Design of scFv-Fcs, mutagenesis, and binding assays

**T**he heavy and light variable chains of Mab14 (43) were joined by a linker sequence of 3 × (Gly, Gly, Gly, Ser) to prepare a single chain variable fragment (scFv), and cloned behind the gp68 signal sequence. This was linked through an additional flexible linker containing a thrombin cleavage site to the Fc portion of human IgG1, and a C-terminal 6-His tag added (35). Residues changed in the antigen binding region of the scFv included those predicted to be directly interacting with the viral capsid, or to be interacting with capsid residues that control specific Mab14 binding, including capsid residue 93 (19). Mutagenesis was conducted through the use of Phusion Site-Directed Mutagenesis Kit (Thermo Fisher Scientific, Waltham, MA). Bacmids were produced through recombination between the DH10Bac (Invitrogen) vector and the pFastbac donor plasmids. Sf9 insect cells (Invitrogen) were then transfected with the bacmids to produce the initial stock of baculovirus. Proteins were expressed from High Five cells over five days incubation at 28°C. The culture supernatant was centrifuged at 10,000 RPM for 30 min before being dialysed into 50 mM Tris-HCl, 150 mM NaCl, 0.05% NaN_3_, (pH 7.5). The scFv-Fc was isolated with a Protein G column, then passed through a Sephacryl S-200 chromatography column in PBS (GE Healthcare, Piscataway, NJ), and the protein in the monomeric protein peak was collected.

The BLItz system (ForteBio, Menlo Park, CA) was used to measure binding kinetics, using Protein A biosensors which were first blocked with kinetics buffer (PBS with 0.01% BSA, 0.02% Tween 20, 0.005% NaN3). Incubations included 300 secs baseline, 300 secs loading, 60 secs baseline, 300 secs association, and 300 secs dissociation. Antibodies were tested at various dilutions, and used at their optimum loading concentration. Viral capsids were added at 0.3 mg/mL. The BLItz Pro software (ForteBio) was used to analyze the data, and statistical analyses of binding assays were conducted with GraphPad Prism 5 (GraphPad Software, Inc., La Jolla, CA). Error bars on **Fig. 5** represent the mean ± SEM obtained through multiple independent experiments.

## RESULTS

### Independent CPV and Fab 14 incubations generated complexes with complete and partial Fab occupancies

Here we examine and compare the results obtained from two separately prepared datasets of Fab 14-CPV complexes. As can be seen in the micrographs (**Fig. 1**), one of the resulting complexes had near-saturation of the Fab binding sites, whereas the other showed only partial Fab occupancy. The complete Fab occupancy, defined as close to 60 Fab 14 molecules bound per capsid, was termed “full-Fab” and allowed icosahedral symmetry averaging to reveal bound Fab. The partial Fab occupancy of binding sites, described as fewer than ~20 Fab 14 molecules per capsid, was called “low-Fab” and was used for an initial icosahedrally averaged reconstruction, that was followed by an asymmetric exploration of the complex and the development of algorithms to assess binding patterns.

**Figure 1.**
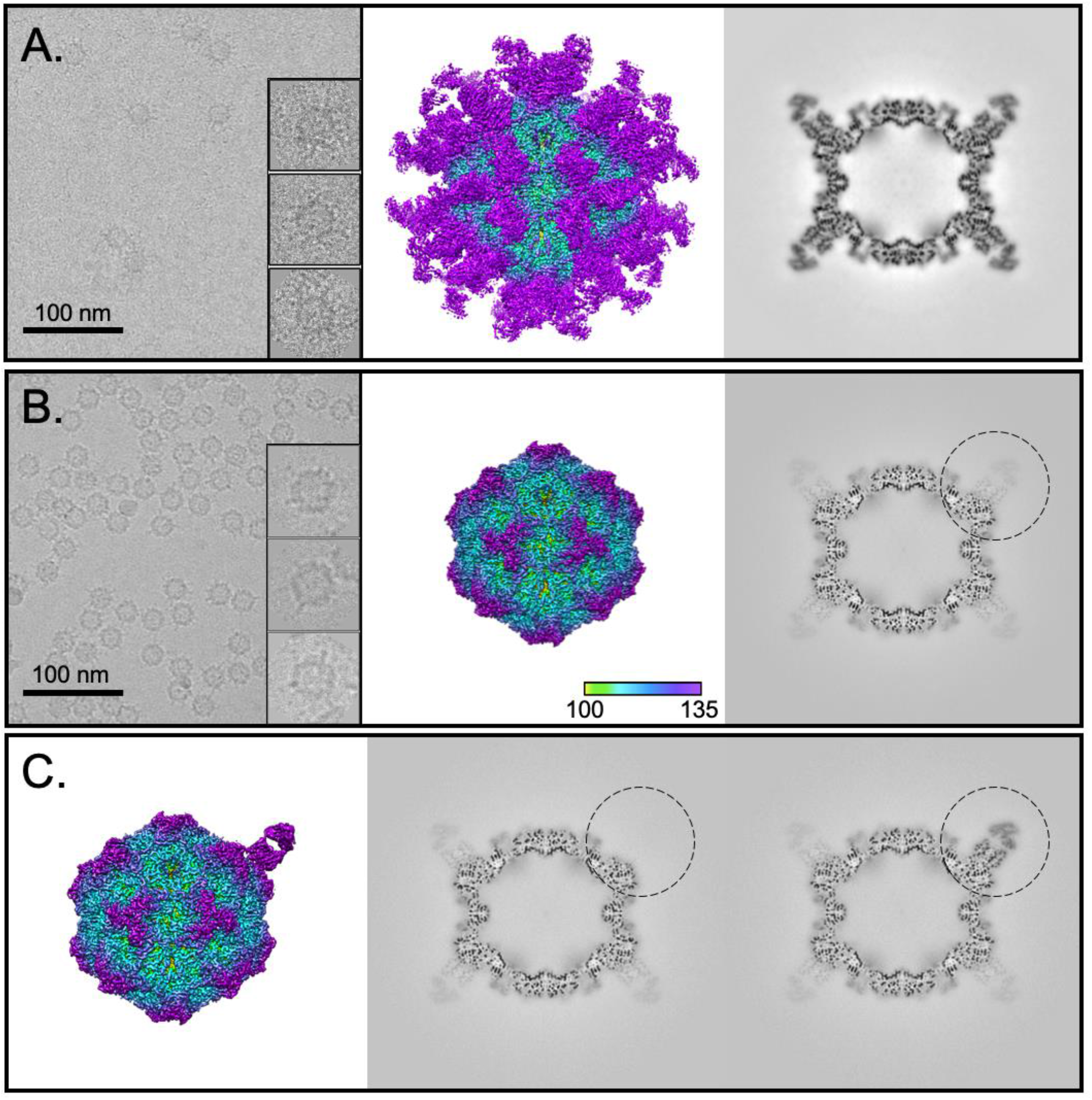
Full-Fab and low-Fab data reconstructed with and without icosahedral symmetry averaging. A and B) (Left) In the cryo-EM micrographs, an excess of Fab in the incubation results in Fab attached to most capsid binding sites, which can be seen as full-Fab complexes that have a spidery appearance. In comparison, complexes resulting from lower ratio of Fab:virus incubation have obvious low fab occupancy. Scale bar is 100nm. (Center and right) The 3.2Å and 2.3Å icosahedrally averaged cryo-EM maps of Fab 14 - CPV full-Fab and low-Fab complexes are surface rendered and colored according to radius, with key inset. The central section of each corresponding map shows the magnitude of Fab density relative to that of the capsid. (C) Asymmetric reconstruction of low-Fab data using localized classification approach. Although there were many more Fab-unoccupied particles per particle, we intentionally selected only as many Fab-unoccupied subparticles as there were Fab-occupied subparticles from each particle. (Left) 2.4Å surface rendered asymmetric map is colored according to radius (as above) and shown with (center and right) the central sections of the fab unoccupied and occupied subparticles (black circle) in the context of the capsid. The low magnitude Fab density seen in other positions in the central section corresponds to an average of the other 10 Fab molecules averaged over the remaining 59 sites.

### Reconstruction of the full-Fab data

Using icosahedral symmetry averaging in cryoSPARC, the full-Fab data were refined to a 3.2 Å resolution map (**Fig. 1 and S.Fig 2**) (27). The crystal structures for Fab 14 (PDB ID 3GK8) (19) and the CPV capsid (PDB ID 2CAS)(17) were fitted into the corresponding densities to initiate the builds followed by manual adjustment in COOT and simulated annealing in PHENIX. The resulting capsid structure superimposed on 2CAS with an RMSD of 0.503 (with C-alpha), with the only significant difference mapping to loop 228 (His 222 - Thr 230) in the virus structure. All other areas that worsened the RMSD were the result of other flexible loop movements during simulated annealing in the weak density of fivefold loop residues 156-162, as well as residues 360-375 and 510-520. The only substantial change between the crystal structures and the cryo-EM density was in the binding interface, where Fab residues 100 and 101 were out of density. This discrepancy corresponds to a polymorphism of Fab 14 residue 100 in the H chain, which was identified as Phe in the original Fab sequence (19); however, a Ser was identified in this position on resequencing of the variable domain (**Table S2**). The density in our structure supports the assignment as Ser 100. After changing the identity of the side chain, Ser 100 and His 101 were adjusted to fit into the Fab density within the complex. With these modifications, no other significant density differences were interpreted in the binding interface, and the fitted Fab crystal structure was used without further refinement for interpretations.

Local resolution mapping showed that the capsid shell had somewhat better resolution than the bound Fab, which likely correlates to slight flexibility. Virus-to-Fab contacts were identified as residues having atoms separated by less than 0.4 Å van der Waal’s radius (36). The newly identified Fab 14 footprint is different than what was estimated previously from the 12 Å resolution capsid-Fab structure (19) (**Table 1**). The main interface interactions took place between capsid surface loops containing residues 93 and 228 that interacted with CDRs H2 and 3 and L1 of the antibody.

**Table 1:**
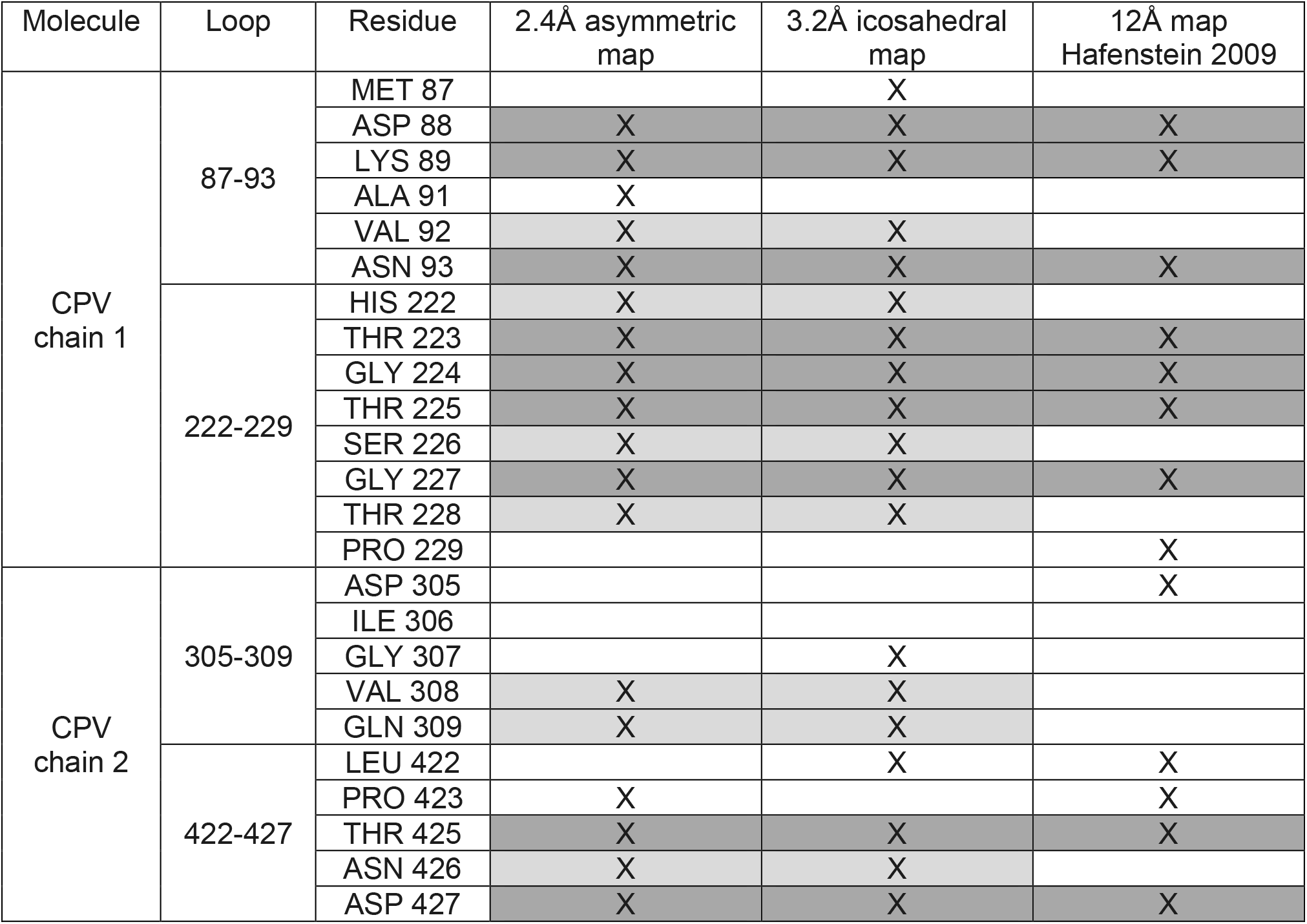
Footprint of Fab 14 on CPV capsid

| Molecule | Loop | Residue | 2.4Å asymmetric map | 3.2Å icosahedral map | 12Å map<br>Hafenstein 2009 |
| --- | --- | --- | --- | --- | --- |
| CPV<br>chain 1 | 87-93 | MET 87 |  | X |  |
|  |  | ASP 88 | X | X | X |
|  |  | LYS 89 | X | X | X |
|  |  | ALA 91 | X |  |  |
|  |  | VAL 92 | X | X |  |
|  |  | ASN 93 | X | X | X |
|  | 222-229 | HIS 222 | X | X |  |
|  |  | THR 223 | X | X | X |
|  |  | GLY 224 | X | X | X |
|  |  | THR 225 | X | X | X |
|  |  | SER 226 | X | X |  |
|  |  | GLY 227 | X | X | X |
|  |  | THR 228 | X | X |  |
|  |  | PRO 229 |  |  | X |
| CPV<br>chain 2 | 305-309 | ASP 305 |  |  | X |
|  |  | ILE 306 |  |  |  |
|  |  | GLY 307 |  | X |  |
|  |  | VAL 308 | X | X |  |
|  |  | GLN 309 | X | X |  |
|  | 422-427 | LEU 422 |  | X | X |
|  |  | PRO 423 | X |  | X |
|  |  | THR 425 | X | X | X |
|  |  | ASN 426 | X | X |  |
|  |  | ASP 427 | X | X | X |

### Reconstruction of the low-Fab data

In an initial step, the low-Fab data were reconstructed while imposing full icosahedral symmetry, which resulted in a 2.3 Å resolution map (**Fig. 1**). As expected, the low occupancy of Fab resulted in weak density due to the 60-fold averaging of the fewer than 60 Fab molecules per capsid. Consequently the Fab structure could not be interpreted. The capsid build was initiated with the fitted crystal structure (PDB ID 2CAS) and further refined in Phenix (17, 29). Unresolved regions included the 226-228 loop where the density was non-continuous, weak, and uninterpretable.

### An asymmetric reconstruction of the low-Fab data resolved the bound Fab and allowed the Fab structure to be built in the context of the virus

Using the icosahedrally-averaged reconstruction of low-Fab data as a starting point, an asymmetric reconstruction was done to resolve the Fab density in the context of the whole capsid. Subparticle extraction and 3D classification were performed to classify subparticles as either Fab-occupied or -unoccupied (**Fig. 1 and S.Fig. 1**), showing that an average of 10 Fab were bound per capsid in the low-Fab data. The results of subparticle classification were used to generate an asymmetric map after superimposing all the Fab density in a standard orientation (ISECC_symbreak), resulting in a final asymmetric map at 2.4 Å global resolution (**Fig. 1 and S.Fig. 2**). In this asymmetric map, the virus-Fab interface had stronger density at higher resolution than the icosahedrally-averaged full-Fab map (**S.Fig. 3**). The Fab 14 crystal structure was fitted to initiate the build with the identity of residue 100 corrected from Phe to Ser, as above, and additional refinement was done in COOT and PHENIX (28, 29).

### Both Fab-occupied and unoccupied virus capsid structures were reconstructed

To compare directly the Fab-occupied and -unoccupied structures, a matched map was generated from the same particles, orienting them to feature no Fab density at that same location (**Fig. 1**). Direct comparability was maintained between the two asymmetric reconstructions, such that individual particles contributed equally to both maps, differing only in their orientations. For example, if particle A was determined to possess eleven Fab molecules, it was incorporated in only eleven orientations in both maps, by discarding the surplus unoccupied sites. This selection criteria ensured that particles contributed equally to both reconstructions, a process we termed “particle matching.” The resulting 2.4 Å resolution asymmetric map resolved the unoccupied Fab-binding site.

### The Fab-occupied and -unoccupied capsid epitopes differ at the 228 loop

To test for local conformational changes induced by bound Fab, a difference map was calculated between the particle-matched asymmetric reconstructions of Fab-bound and -unbound epitopes (**Fig. 2a**). In addition to the expected Fab difference density, there was significant capsid difference density corresponding to virus loop 228 in the Fab binding interface. As described above, the fitted crystal structure of the capsid protein (PDB ID 2CAS) was refined in COOT and PHENIX for the Fab-bound and -unbound maps (**Table S3**) (17). Inspection of the bound- and un-bound models suggests a hinge-like motion of the 228 loop (1.9 Å maximum Cα-Cα displacement), which makes room for Fab heavy chain residue His101 (**Fig. 2b and S.Movie 1, 2**). The bound and unbound structures superimposed with a Cα RMSD of 0.200Å, indicating minor variation between the two structures due to reconfiguration of residues 226-229 in the Fab interface.

**Figure 2.**
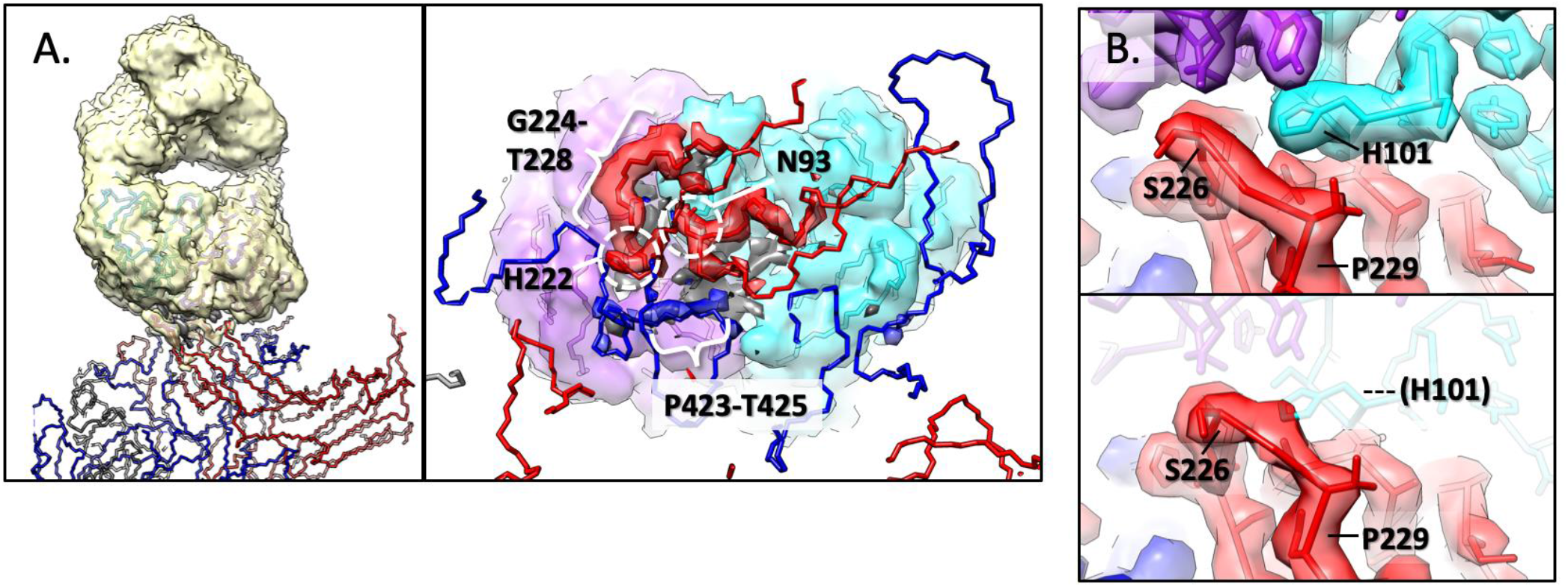
Fab-virus interface and interactions. (A, Left) Difference mapping showing all positive density (taupe) most of which corresponds to Fab14; however, there is additional positive (taupe) and negative (dark grey) density located at the Fab-capsid interface. Two copies of the VP2 capsid protein (red and blue wire) comprise the binding site. (Right) Map of interface is rotated 90° with the virus slabbed to show the interface and Fab heavy (cyan) and light chains (purple) surface rendered. This zoomed view illustrates positive difference density (red and blue) corresponding to the virus loops. In the absence of Fab some virus density is discontinuous and cannot be built (dashed lines). The positive difference density corresponds to capsid residues 88-94, 222, 224-228, and 423-425 (colored by nearest chain). Negative difference density (dark grey) can be seen adjacent to the positive difference density. (B) Zoomed image of the virus (red and blue) and Fab (purple and cyan) interface. The VP2 228 loop is displaced by a maximum of 1.9Å upon interaction with Fab heavy chain residue H101 (top vs bottom). This same view is provided as a morph-map in S.Movie 2.

### The Fab footprint identified using the asymmetric map is nearly the same as the footprint identified using the icosahedrally averaged map

After Fab and capsid structural refinement, residues in the interface were defined as contacts using the same method as with the full-Fab icosahedrally averaged map (**Table 1**). Predicted contacts in the 222-229 loop were identical in both the 3.2 Å and 2.4 Å resolution maps. For the 87-93, 305-309, and 422-427 loops, the improved resolution of the asymmetric map moved a few residues (Met87, Ala91, Gly307, Leu 422, Pro423) either in or out if the strict contact criteria cutoff (**Table 1, Fig. 3**). Virus capsid residues 93, 222, and 224 identified in the footprint have also been shown previously to influence Mab14 binding and identified as selected escape mutations (11, 12, 15, 16). Local resolution mapping showed small improvements in resolution of the surface epitope upon Fab binding, including residues 423-426, 88-93, and 309 (**Fig. 4**). Residues not involved in direct Fab interaction had similar local resolution in the two maps, suggesting that Fab-binding stabilizes loop in the epitope by interacting with and burying the capsid surface.

**Figure 3.**
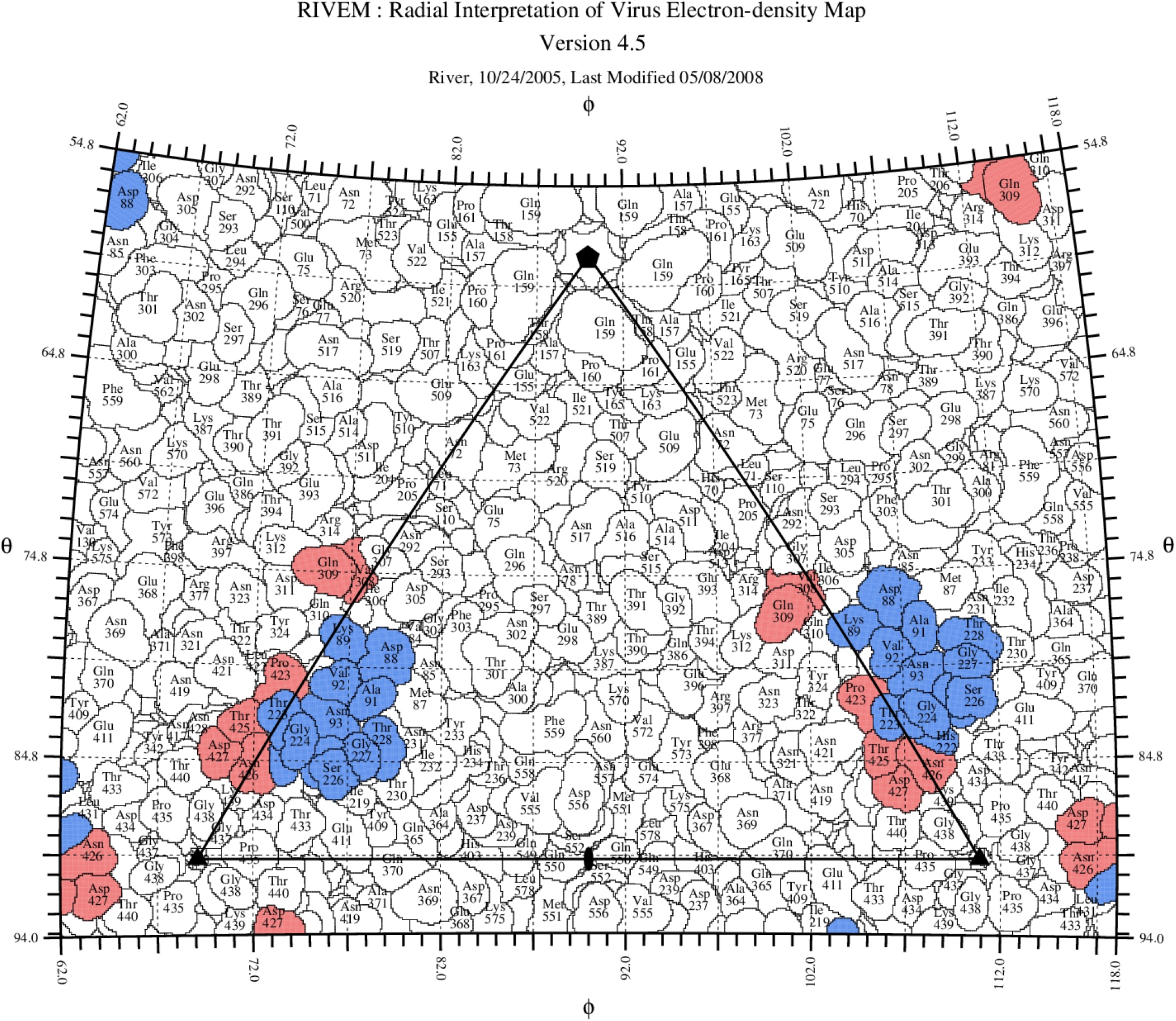
Road map of the Fab 14 footprint on the CPV surface. The capsid surface is shown as a stereographic projection where the polar angles phi and theta represent the latitude and longitude of a point on the virus surface (46). The virus surface is represented as a quilt of amino acids (47), and the icosahedral asymmetric unit of the virus is indicated by the triangular boundary. The footprint of Fab 14 has contributions from symmetry-related copies of the capsid protein (red and blue) as in Fig. 2.

**Figure 4.**
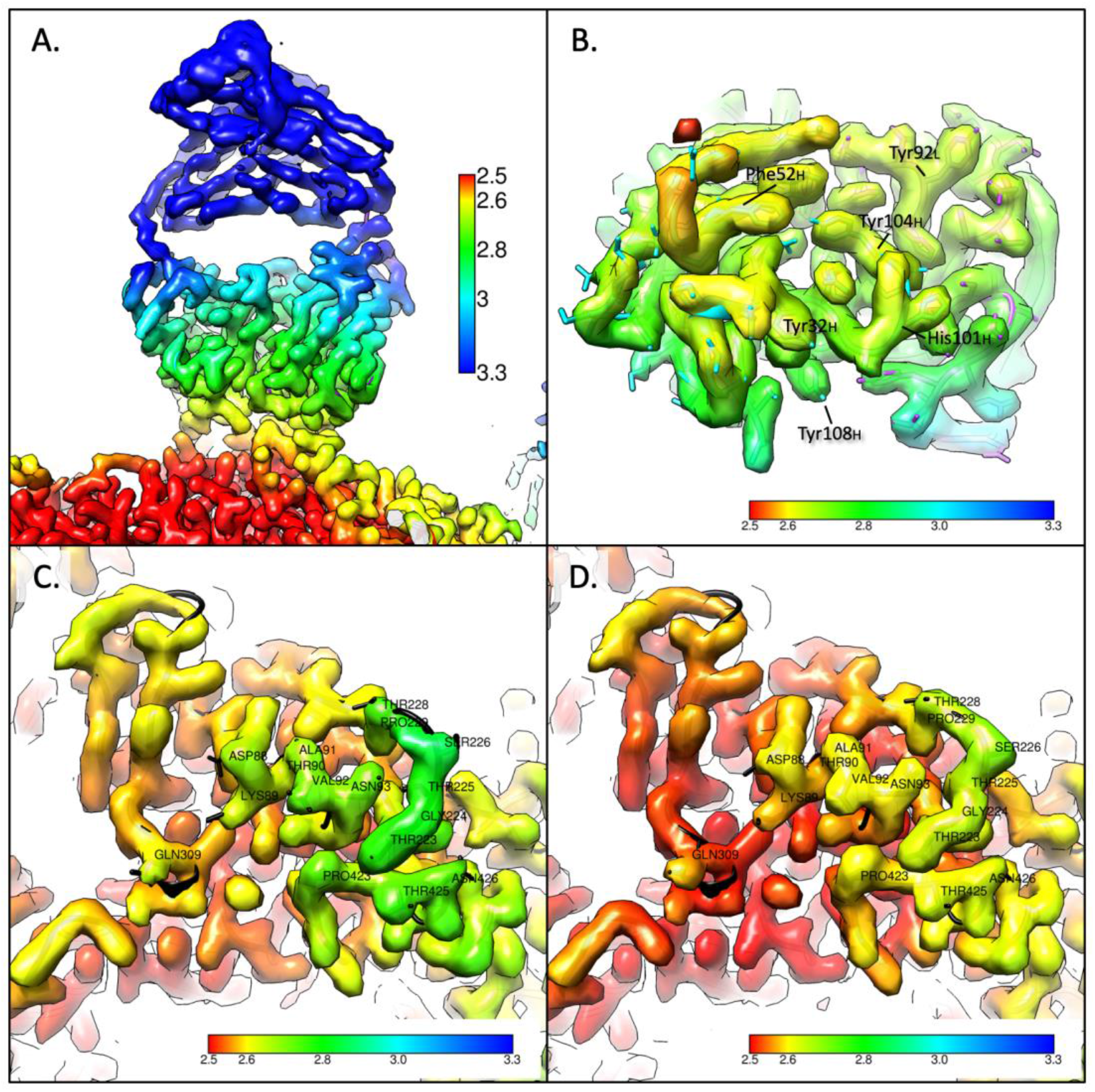
Local resolution maps colored according to the resolution shown in the key in the first panel. A) The surface rendered map for the Fab reconstructed by subparticle classification shows that distal portions of the Fab have poorer resolution, likely due to flexibility of the elbow. B) The Fab map rotated 90° to show the region of contact with the virus has consistently good resolution of 2.6-2.8 Å. C) Region of contact on the virus capsid for unoccupied and occupied Fab binding illustrate that the Fab binding orders the density in the 228 loop. The overall resolution of the virus contact region is improved upon Fab binding suggesting the stabilization of the epitope.

### Targeted mutations inhibit Fab 14 binding

Mutations of the virus that influenced Mab14 binding were identified in previous studies as residues 93, 222, and 224 (11, 12, 15, 16). Specifically a change in Asn93 abrogated binding as demonstrated by the diminished ability of FPV (Lys 93) to bind Mab 14. Mutations G224R, G224E, and H222Y have also been known to interfere with the Mab14 binding. For further testing, the Fab 14 was expressed as an scFv-Fc, which allowed us to examine its binding to the capsid, and to create mutations in selected CDR loops. The scFv-Fc bound to CPV capsids to a higher level than to FPV capsids (**Fig. 5**), as expected due to the known specificity of the virus for CPV (11, 16). Mutations in two different loops within the complementarity determining regions (CDRs) of the antibody resulted in greatly reduced binding to CPV (and even less binding to FPV), confirming that those were involved in critical contacts in the binding of the antibody to the capsid (**Fig. 5**).

**Figure 5.**
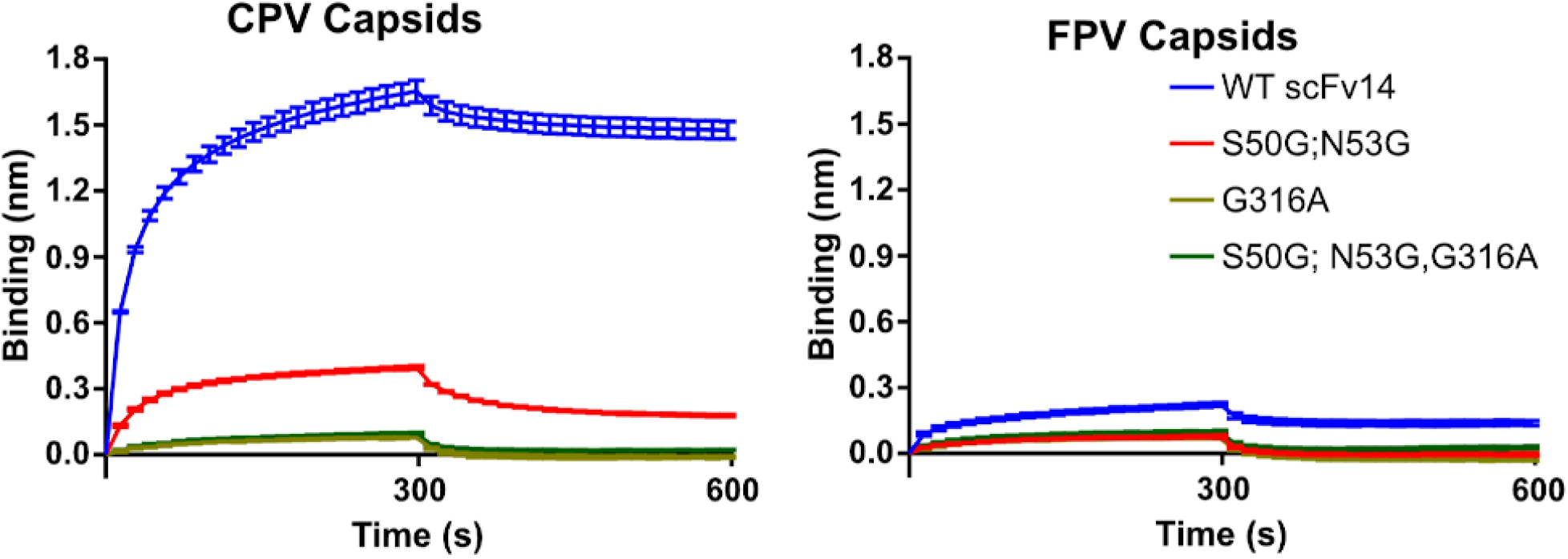
Blitz binding data. Levels of binding to (A) CPV or (B) FPV capsids of wildtype scFv of Fab 14 or scFv14 variants with mutations in the complementarity determining region. Protein A biolayer interferometry probes were incubated with scFv-Fc to the same level of bound protein, washed, incubated with CPV-2 or FPV empty capsids, then incubated in buffer. Mutant scFv14 forms were Ser50Gly and Asn53Gly together, Gly316Ala alone, or all three mutations.

To predict the effect of the FPV-encoded Lys at position 93 on the antigenic epitope, the FPV and CPV crystal structures were fitted into the Fab-bound density map for comparison. The longer side chain of Lys 93 in FPV followed the same trajectory as that of Asn 93 of CPV, but extended out of cryo-EM density, and has been predicted to form two hydrogen bonds to the carbonyl oxygen atoms of residues Thr 225 and Gly 227 (37). The 228 loop is immediately adjacent to residue 93 (**Fig. 4c-d**). In the crystal structures of CPV and FPV that were not antibody bound, the loop containing residue 228 extended out of density when compared to that seen in the Fab-bound structure, consistent with the ordering of the 228 loop due to Fab binding.

Other differences in the Fab binding interface of CPV compared to FPV were investigated by mapping the electrostatic surface potentials to understand how the Lys (FPV) or Asn (CPV) at residue 93 might alter the surface charge. There was positive charge at the Fab surface where it interacts with virus residue 93, consistent with a favorable CPV and impaired FPV interaction, due to the positively-charged Lys 93 of FPV (**Fig. 6**).

**Figure 6.**
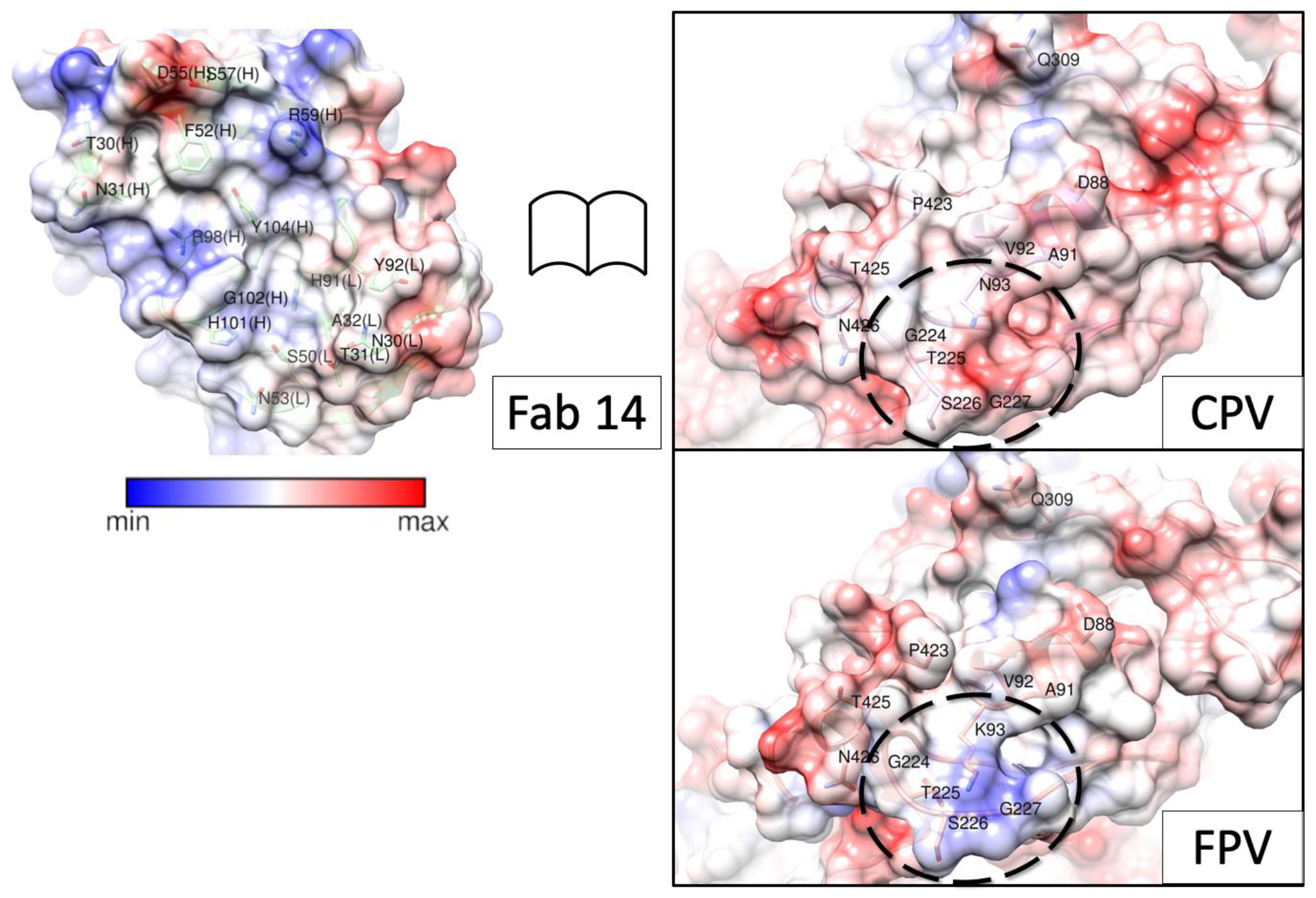
Open book view of the binding interface. Surface rendered maps depicted with columbic potential (key indicates blue is positive and red is negative charge) for Fab and CPV illustrate the complementarity of the binding surfaces outlined by dashed lines on virus capsid surface. For comparison, the same region of FPV was rendered and the change at residue 93 can be seen to alter the surface charge within the region that comprises the epitope.

### Correlated local classification provides a Fab fingerprint of the complex

For the low-Fab dataset, each individual complex was analyzed to identify which of the 60 binding sites were occupied by Fab relative to each other on the same virus capsid. After extraction and classification of the originally defined subparticle (see above), the Fab-occupied particles were identified per complex. Each complex was defined according to the 3D binding of Fab molecules relative to each other in the context of the capsid, which we termed “Fab fingerprint” (**Fig. 7**). This fingerprint is effectively a binarized version of a radial distribution function (RDF), consisting of the center-of-mass distances between all Fab-occupied subparticles. These per-particle Fab fingerprints were pooled and normalized against a hypothetical fully saturated particle. The normalized RDF plots can reveal deviation of the dataset from expected, random-chance binding patterns.

**Figure 7.**
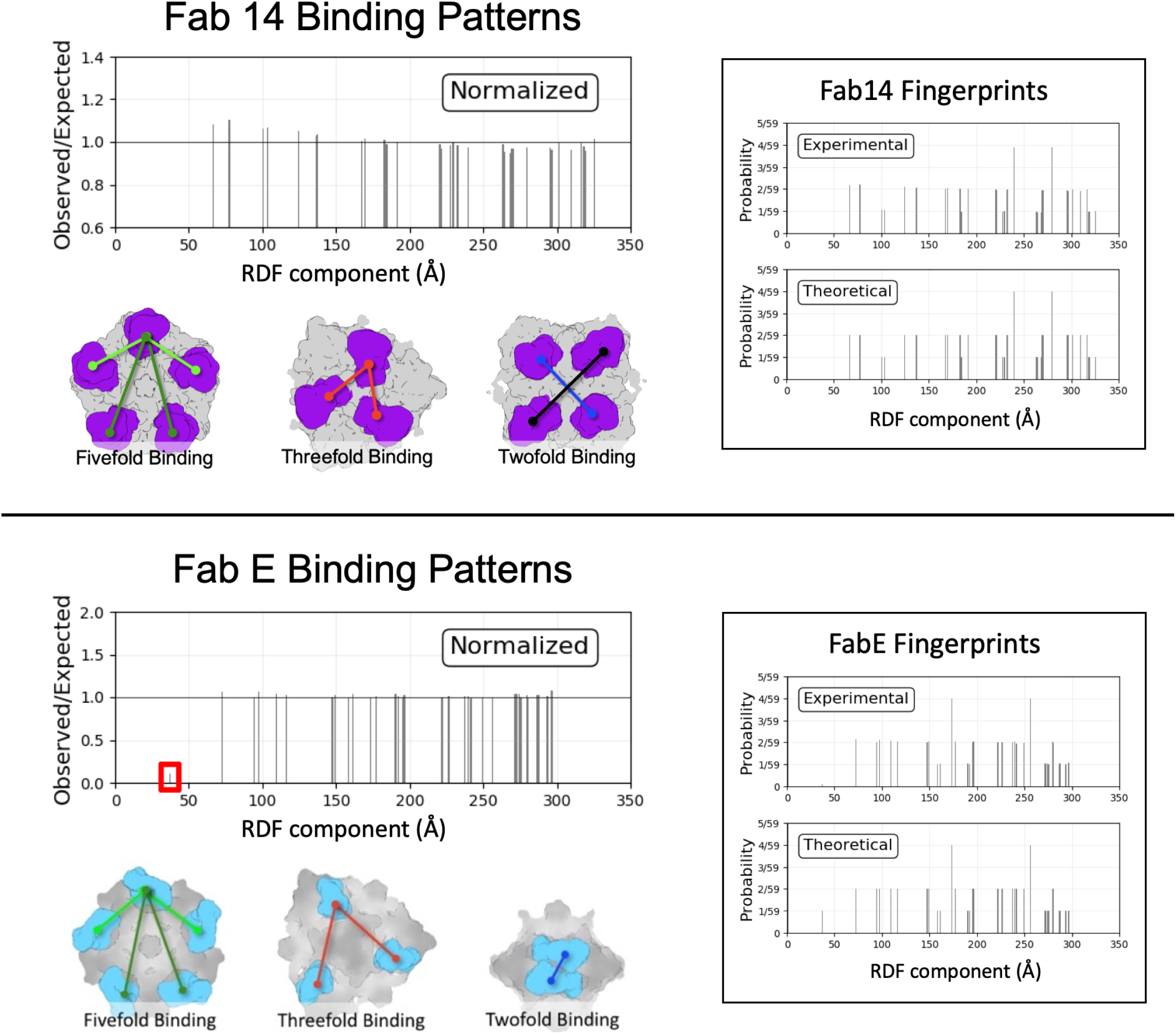
Icosahedral Subparticle Extraction and Correlated Classification (ISECC). Binding Patterns for Fab 14 and Fab E are described as Radial Distribution Functions evaluated at binding sites. For Fab 14 (top panel), a modest excess of proximal RDF components was detected after normalization against a theoretical particle with unbiased binding. For Fab E, a severe deficit of the most proximal RDF component was observed, consistent with twofold symmetry clash of Fab E. Example geometry corresponding to RDF components are shown in each case.

Over the entire low-Fab dataset, Fab 14 binding patterns largely matched random-chance predictions, but with a slight excess of the proximal RDF components, corresponding to increased binding events among nearby sites (**Fig. 7, upper panel**). Thus, our analysis suggested that there might be binding cooperativity of Fab 14 molecules at the local level, since they tend to cluster on the capsid rather than disperse randomly across the 60 symmetry-related epitopes.

For comparison we used the same correlated local classification method to calculate fingerprints for another Fab-CPV complex. That previously determined cryo-EM structure showed that the Fab of monoclonal antibody E (Fab E) recognized a different epitope on the capsid located close to the two-fold axis, with moderate clash between symmetry-related Fabs (38). The virus-Fab E map central section revealed a magnitude of Fab E density less than half that of the capsid itself. As expected, correlated local classification showed a strong deficit of the most proximal RDF component, consistent with steric clash of Fab E molecules across the twofold symmetry axis. All other RDF components matched expected values **(Fig. 7, lower panel)**. This negative cooperativity contrasts with the modest positive local cooperativity observed with Fab 14. Residual signal from the twofold Fab E RDF component may be due to low levels of classification error in the modest resolution dataset.

To further validate RDFs as a tool for correlated classification, two undersaturated enterovirus-Fab complexes were analyzed, one without any steric clashes induced by Fab binding, and the other complex featuring strong steric collisions about each threefold symmetry axis (**S.Fig. 4**). The behavior of CPV-Fab14 matched that of the clash-less enterovirus-Fab complex. Likewise, the behavior of CPV-FabE matched that of the enterovirus-Fab complex with the threefold symmetry clash. These results establish correlated classification as a tool for detecting and quantifying cooperativity of Fab binding.

## DISCUSSION

The small and seemingly simple parvovirus capsid performs many different functions in the process of packaging the ssDNA genome, trafficking between cells and animals, and entry into a permissive host. Critical events include effective receptor binding, endocytosis and transport of the genome to the nucleus, as well as binding and potential evasion of the host antibody response. Interactions with antibodies require recognition and attachment to epitopes displayed on the assembled virus capsid, which may lead to neutralization of infection or select for antigenic variation of the virus. The relationships between antibody binding sites and evolution that results in antibody escape, are complicated by the overlap with receptor binding sites needed for cell infection. Here we used multiple approaches to analyze the Mab14 that binds to the CPV capsid at a position that also controls TfR recognition and the viral host ranges. The results of this study explained the specificity of binding to CPV compared to FPV and also revealed Fab binding patterns.

The traditional structural approach to reveal a specific interaction on the surface of an icosahedral virus is to saturate all the potential binding sites on the capsid by incubating purified particles with excess protein. This forced symmetry approach often results in a virus-protein complex that is coated with protein and allows for icosahedral symmetry averaging during the reconstruction process. The averaging often allows higher resolution to be achieved, but various factors may limit success, including low affinity and steric collisions of symmetry-related bound molecules or bulky proteins. However, complexes made with full occupancy of host protein binding sites on icosahedral viruses may poorly represent *in vivo* virus host interactions, which are more often asymmetric or heterogeneous.

Using asymmetric reconstruction and innovative subparticle classification techniques, we were able to use a single data set to solve the structures of Fab-bound and unbound epitopes within the same complex. The local conformational changes to the virus capsid which may be induced by Fab binding (**Fig. 2, 4, S.Movie 1, 2**) might contribute to host range or tropism, along with the difference in surface charge displayed by the Asn (CPV) and Lys (FPV) at VP2 residue 93. At the local resolution of this region (~2.8 Å) movement of the 228 loop by 1.9 Å is not definitive, however this loop shows higher B factors in the crystal structures of CPV than FPV, supporting flexibility (PDB IDs 2CAS, 1C8D, 4DPV)(17, 18, 39). Host range specificity for CPV and FPV is controlled by residues 93 and 323 of the major capsid protein (16) that together allow specific binding of canine TfR, which is not recognized by FPV (40).

### Antibody binding and specific recognition

As expected, the footprint revealed by our atomic resolution model differs from that estimated previously from a 12 Å resolution structure (19), and we now have an unambiguous definition of the interaction site. The movement or ordering of the residues comprising loop 228, which may be essential for high affinity Fab 14 binding, appears to represent a case of induced fit resulting from the binding, as has also been seen in many other antibody binding interactions (41–43). Other host protein binding events that induce local conformational changes in virus capsids may have been missed due to structural approaches that have focused on making complexes by fully saturating all potential binding sites and imposing icosahedral symmetry averaging during the reconstruction process.

The structures derived from the Fab-bound and unbound asymmetric maps also suggest a model for how the sidechain of residue 93 controls species-specific binding. Compared to CPV (PDB ID 2CAS), the FPV loop 228 (PDB ID 1C8F) is positioned further away from the main part of the conformational epitope and also stabilized due to the Lys 93 sidechain and the new bonds that are formed with the carbonyl oxygen atoms of residues Thr 225 and Gly 227, and those effects would clearly reduce Fab 14 recognition of FPV. In addition, local charge differences may play a role in controlling specific antibody binding due to the positively charged patch on Fab 14 that correlates to the interaction with virus residue 93. The previously mapped escape mutations such as His 222 also support this model overall, due to interactions with the 426 loop of the neighboring capsid protein subunit, which itself possesses two residues involved in Fab binding (Thr 425, Asn 426). Mutation of Fab-buried residue Gly 224 likely directly interferes with Fab binding via steric interference. Thus the identified mutations may directly block Fab binding or they may alter the configuration of loops within the Fab footprint. We also confirmed key interactions associated with the antibody structure by mutating two CDR loops and expressing the protein as scFvs linked to the Fc of human IgG1. Those changes in light chain CDR2 (Ser50Gly and Asn53Gly) and heavy chain CDR3 (Gly316Ala) reduced the binding to both CPV and FPV capsids.

### The local classification and correlation approach revealed additional details compared to simple icosahedral averaging

Besides the structural solution, we used the low-Fab dataset to reveal positive or negative cooperativity in the binding of two different Fabs that bound to different capsid positions (**Fig. 7**). The use of local classification was an innovative and powerful approach to analyzing the data, and may have many uses in analysis of these and other symmetrical virus structures. Implementing an RDF-style analysis of the low-Fab complexes allowed interpretation of Fab binding patterns, conclusively establishing steric collision in cases of suspected atomic clash, and suggesting positive cooperativity where symmetry clash is not a factor. This also suggests value in purposely obtaining undersaturated occupancies of host protein molecules on a virus particle, to allow the direct comparison of occupied and unoccupied binding sites.

Importantly, correlated local classification is dependent on the accuracy of classification results. Negative local cooperativity due to steric clash is seen in the deficit of proximal components in the RDF plots, whereas positive cooperativity is suggested by an excess of proximal components. However, both situations exist in the presence of systematic classification error, and become compelling with the addition of supporting data, repeatability, and comparison with different virus-Fab complexes. Analysis of additional complexes should reveal common patterns of binding behavior, helping establish the threshold for distinguishing true biophysical phenomena from low-level classification error.

Overall this study confirmed that the binding of an antibody with a capsid is a complex process involving multiple loops of the antibody and virus. Capsid changes that have been selected as host range mutations also alter antibody binding, confirming that single point mutations on either the capsid or antibody are sufficient to prevent attachment (16, 44, 45). We were also able to develop innovative approaches to analyze partially occupied capsids and understand the patterns of Fab binding using a cryo-EM structure.

## ACKNOWLEDEGMENTS

Funding was provided by the Pennsylvania Department of Health CURE funds. Research reported in this publication was supported by the Office of the Director, National Institutes of Health, under Award Numbers S10OD019995 (JFC) and S10RR031780 (SH), as well as NIH grants R01AI107121 (SH), R01AI092571 (CRP), and T32CA060395 (LJO).

## Accession numbers

The cryo-EM maps of the icosahedrally refined Low-fab and Full-fab CPV complexes, and asymmetric Fab-occupied and -unoccupied maps are deposited in the EM data bank (www.emdatabank.org/) under accession numbers EMD-XXXX, XXXX, XXXX, and XXXX. Models built for the fab are deposited in the PDB under ID YYYY and YYYY respectively, and models built for the fab-unoccupied and -occupied virus are deposited under YYYY, and YYYY respectively.

## Supplementary Information

**Table S1.** Cryo-EM statistics

| Fab Occupancy | Symmetry | Microscope /Detector | No. of micrographs | Defocus range (u.m.) | No. of particles selected | No. of particles used | Resolution (Å) |
| --- | --- | --- | --- | --- | --- | --- | --- |
| Full-Fab<br>60 Fab/capsid | I2 | FEI Polara, Falcon 3EC | 21,075 | 0.5 – 4.2 | 142,618 | 95,353 | 3.2 |
| Low-Fab<br>10 Fab/capsid | I2 | Titan Krios, Falcon 3EC | 2,265 | 0.7 – 4.9 | 316,808 | 162,627 | 2.3 |
|  | C1/<br>Symmetry<br>-mismatch | Titan Krios, Falcon 3EC | 2,265 | 0.7 – 4.9 | 316,808 | 162,627<br>* 10.2<br>orientations/particle | 2.4 |

**Table S2.**
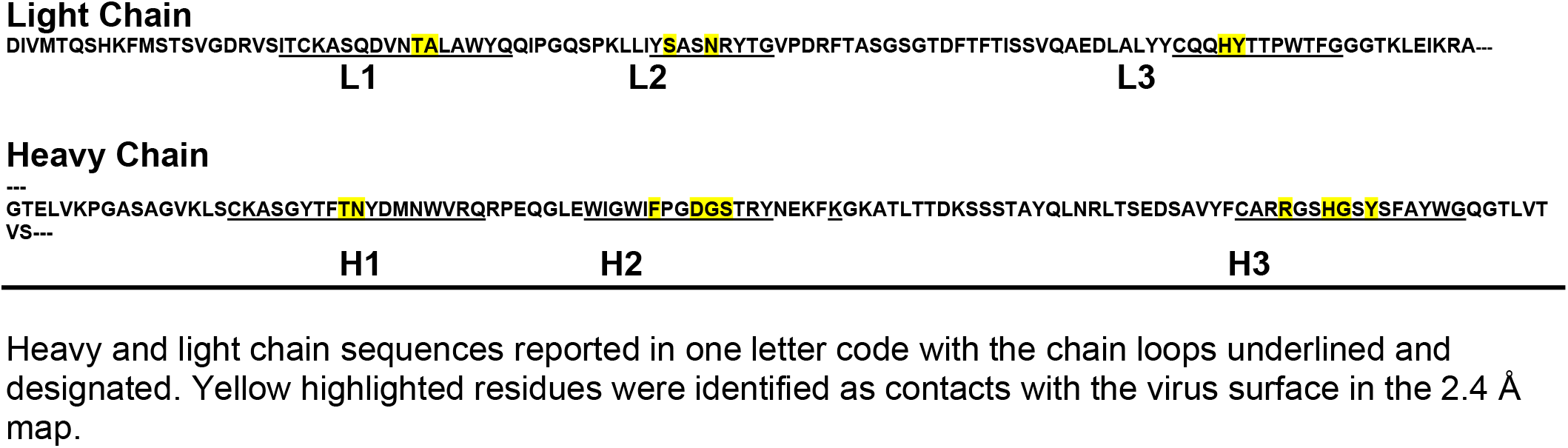
Sequence of the Fab and ScFv

**Table S3:**
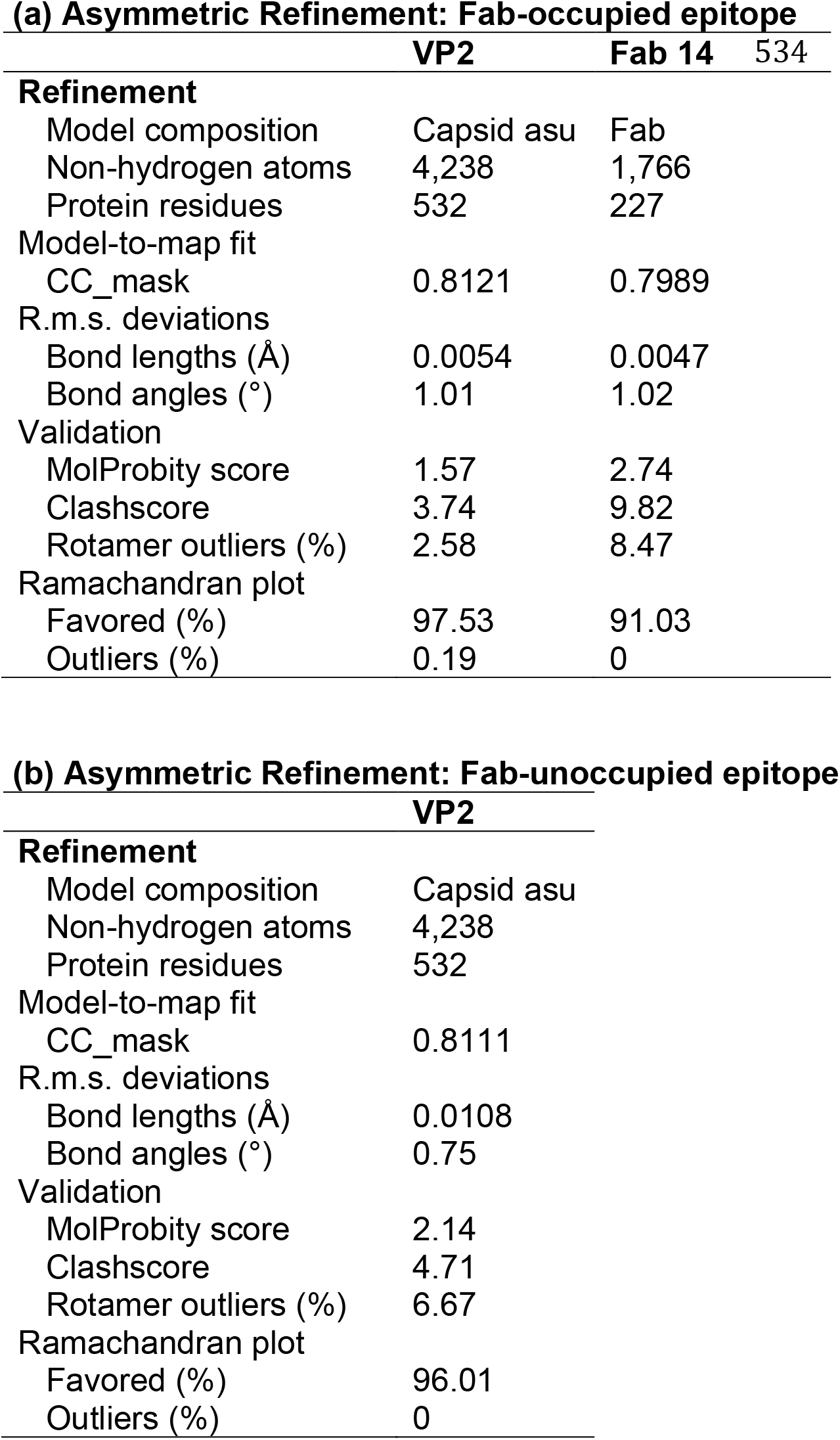
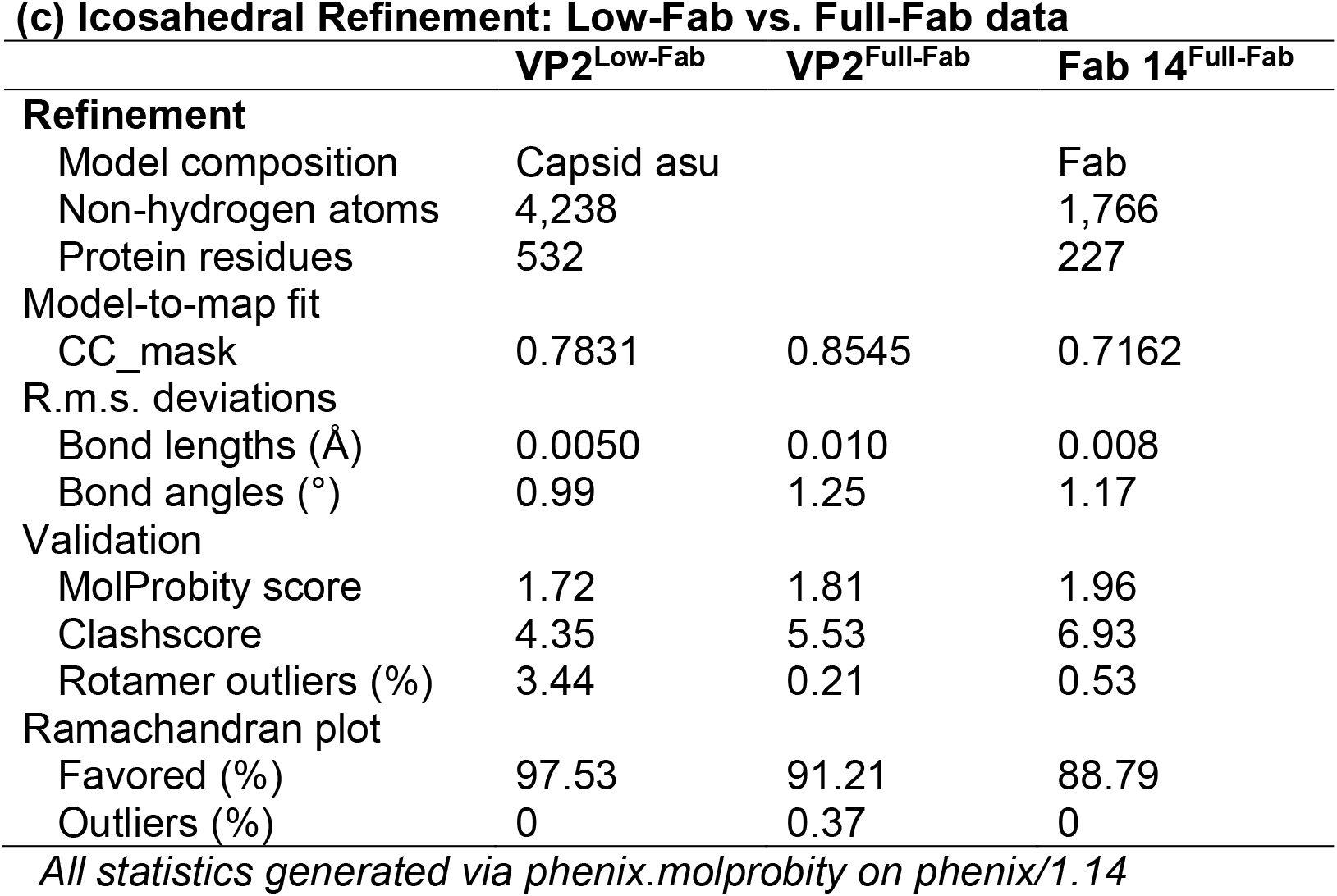
Refinement and Validation Statistics

|  |  |  |  |
| --- | --- | --- | --- |
| 532 | **Table S3: Refinement and Validation Statistics** |  |  |
| 533 | **(a) Asymmetric Refinement: Fab-occupied epitope** |  |  |
|  |  | **VP2** | **Fab 14** |
|  | **Refinement** |  |  |
|  | Model composition | Capsid asu | Fab |
|  | Non-hydrogen atoms | 4,238 | 1,766 |
|  | Protein residues | 532 | 227 |
|  | Model-to-map fit |  |  |
|  | CC_mask | 0.8121 | 0.7989 |
|  | R.m.s. deviations |  |  |
|  | Bond lengths (Å) | 0.0054 | 0.0047 |
|  | Bond angles (°) | 1.01 | 1.02 |
|  | Validation |  |  |
|  | MolProbity score | 1.57 | 2.74 |
|  | Clashscore | 3.74 | 9.82 |
|  | Rotamer outliers (%) | 2.58 | 8.47 |
|  | Ramachandran plot |  |  |
|  | Favored (%) | 97.53 | 91.03 |
|  | Outliers (%) | 0.19 | 0 |

|  |  |  |
| --- | --- | --- |
| 535 | **(b) Asymmetric Refinement: Fab-unoccupied epitope** |  |
| 536 |  | **VP2** |
| 537 | **Refinement** |  |
|  | Model composition | Capsid asu |
|  | Non-hydrogen atoms | 4,238 |
|  | Protein residues | 532 |
|  | Model-to-map fit |  |
|  | CC_mask | 0.8111 |
|  | R.m.s. deviations |  |
|  | Bond lengths (Å) | 0.0108 |
|  | Bond angles (°) | 0.75 |
|  | Validation |  |
|  | MolProbity score | 2.14 |
|  | Clashscore | 4.71 |
|  | Rotamer outliers (%) | 6.67 |
|  | Ramachandran plot |  |
|  | Favored (%) | 96.01 |
|  | Outliers (%) | 0 |

**(c) Icosahedral Refinement: Low-Fab vs. Full-Fab data**
|  | **VP2<sup>Low-Fab</sup>** | **VP2<sup>Full-Fab</sup>** | **Fab 14<sup>Full-Fab</sup>** |
| --- | --- | --- | --- |
| **Refinement** |  |  |  |
| Model composition | Capsid asu |  | Fab |
| Non-hydrogen atoms | 4,238 |  | 1,766 |
| Protein residues | 532 |  | 227 |
| Model-to-map fit |  |  |  |
| CC_mask | 0.7831 | 0.8545 | 0.7162 |
| R.m.s. deviations |  |  |  |
| Bond lengths (Å) | 0.0050 | 0.010 | 0.008 |
| Bond angles (°) | 0.99 | 1.25 | 1.17 |
| Validation |  |  |  |
| MolProbity score | 1.72 | 1.81 | 1.96 |
| Clashscore | 4.35 | 5.53 | 6.93 |
| Rotamer outliers (%) | 3.44 | 0.21 | 0.53 |
| Ramachandran plot |  |  |  |
| Favored (%) | 97.53 | 91.21 | 88.79 |
| Outliers (%) | 0 | 0.37 | 0 |

*All statistics generated via phenix.molprobity on phenix/1.14*

**S.Figure 1.**
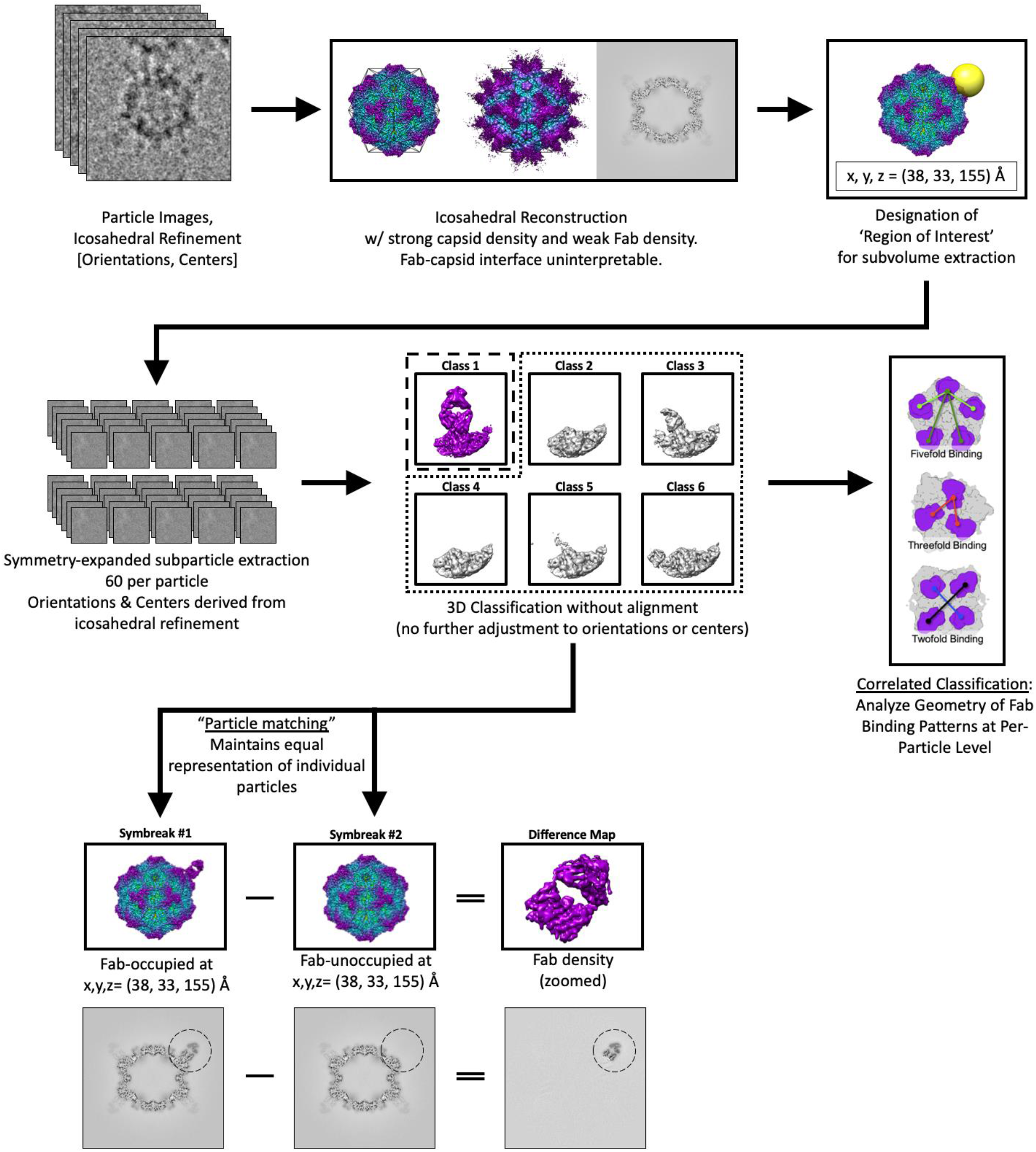
Flow chart of the reconstruction and classification process.

**S.Figure 2.**
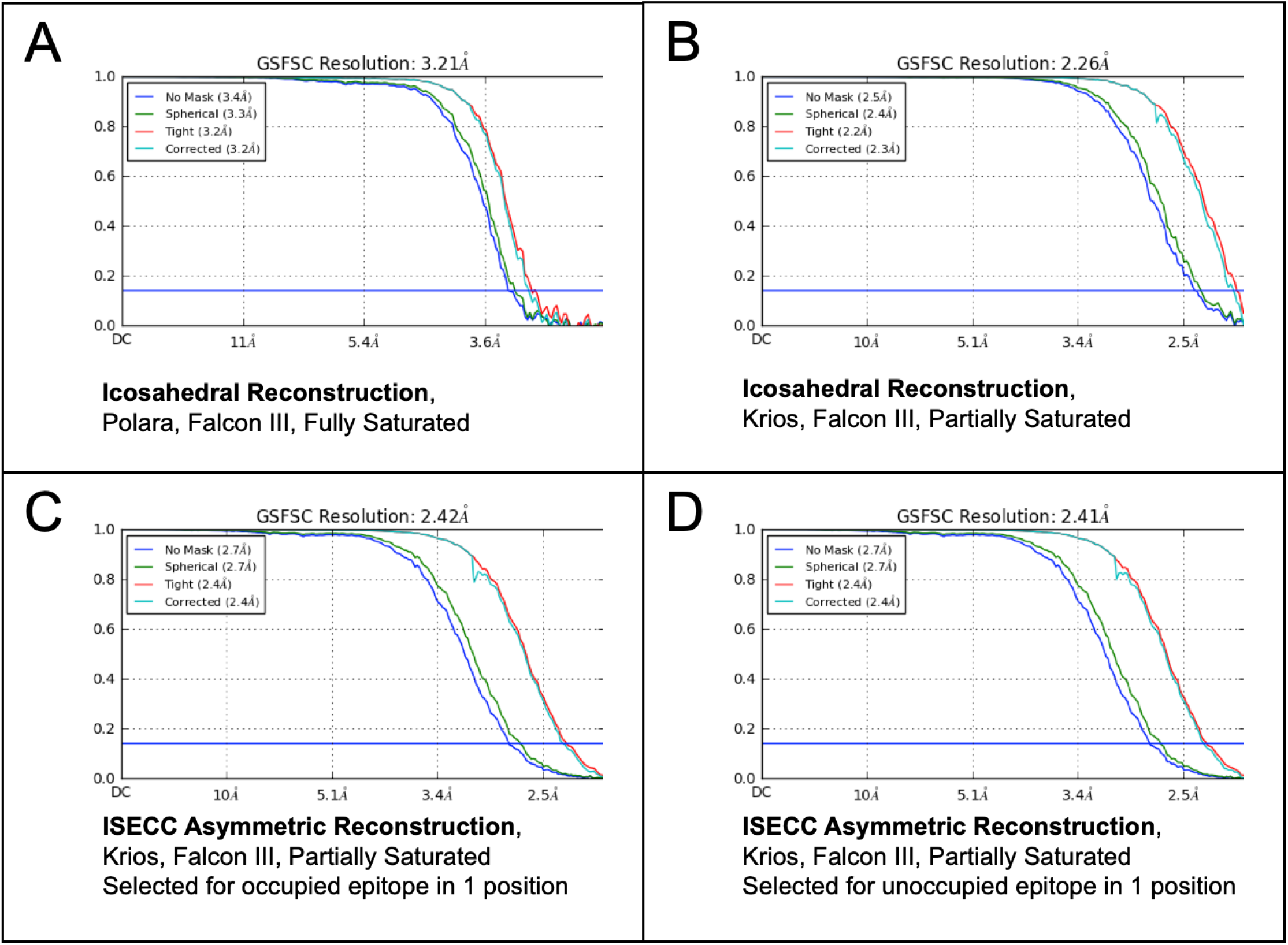
Global FSC curves. **(A, B)** Different instrumentation and sample preparations allowed the partially saturated dataset to achieve higher resolution than the fully saturated dataset (2.3Å vs 3.2Å). **(C,D)** Asymmetric symmetry-break operations sacrificed some of this resolution to resolve the capsid the resolve a selected site on the capsid in the presence or absence of Fab (both 2.4Å global resolution). Local resolution at the selected epitope is shown in Fig. 4.

**S.Figure 3.**
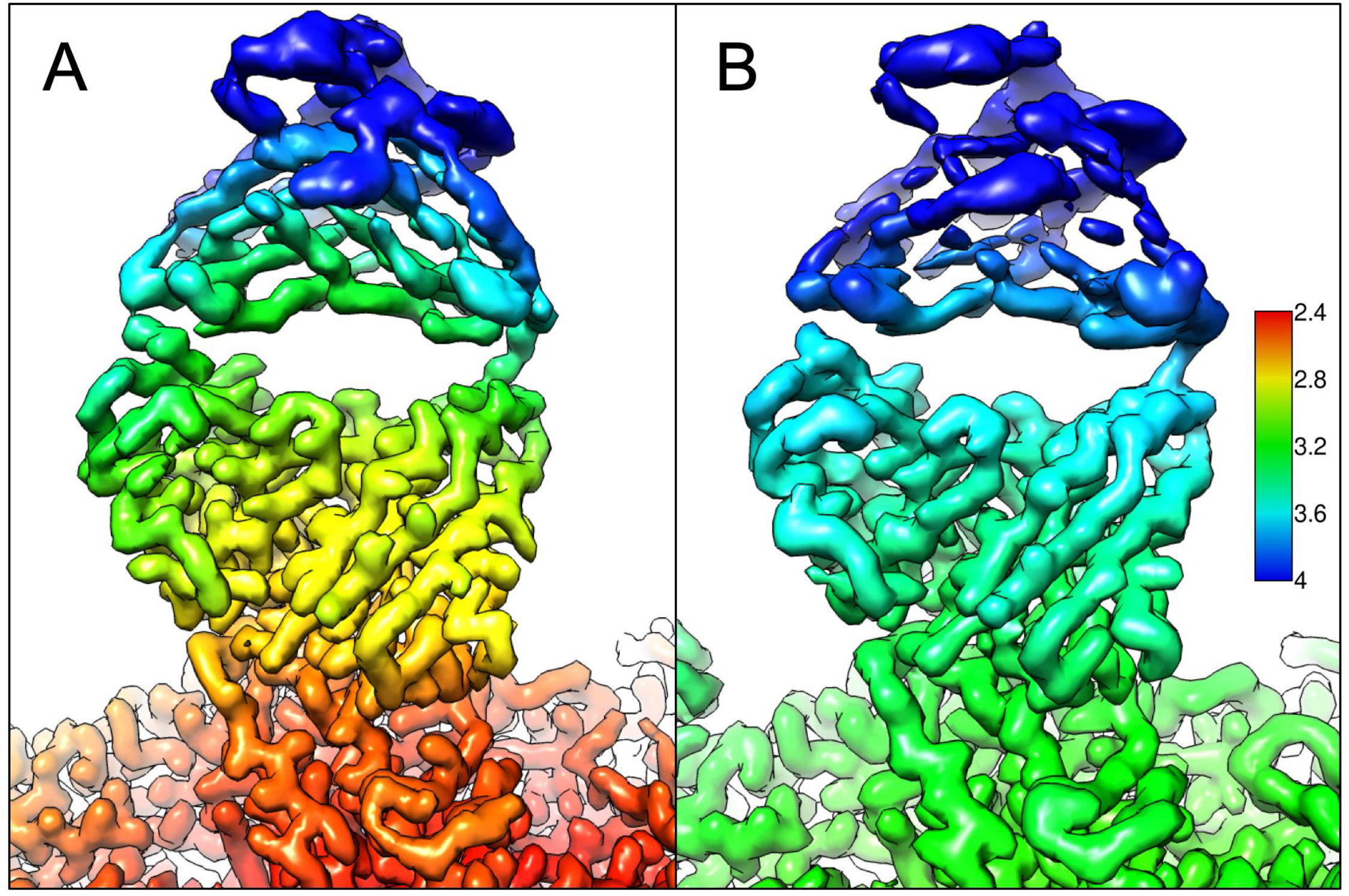
Comparison of whole Fab structures. Fab-density of the asymmetric low-Fab map colored by local resolution (A, left) was consistently better at the Fab-capsid interface than that of the full-fab icosahedrally averaged map (B, left).

**S.Figure 4.**
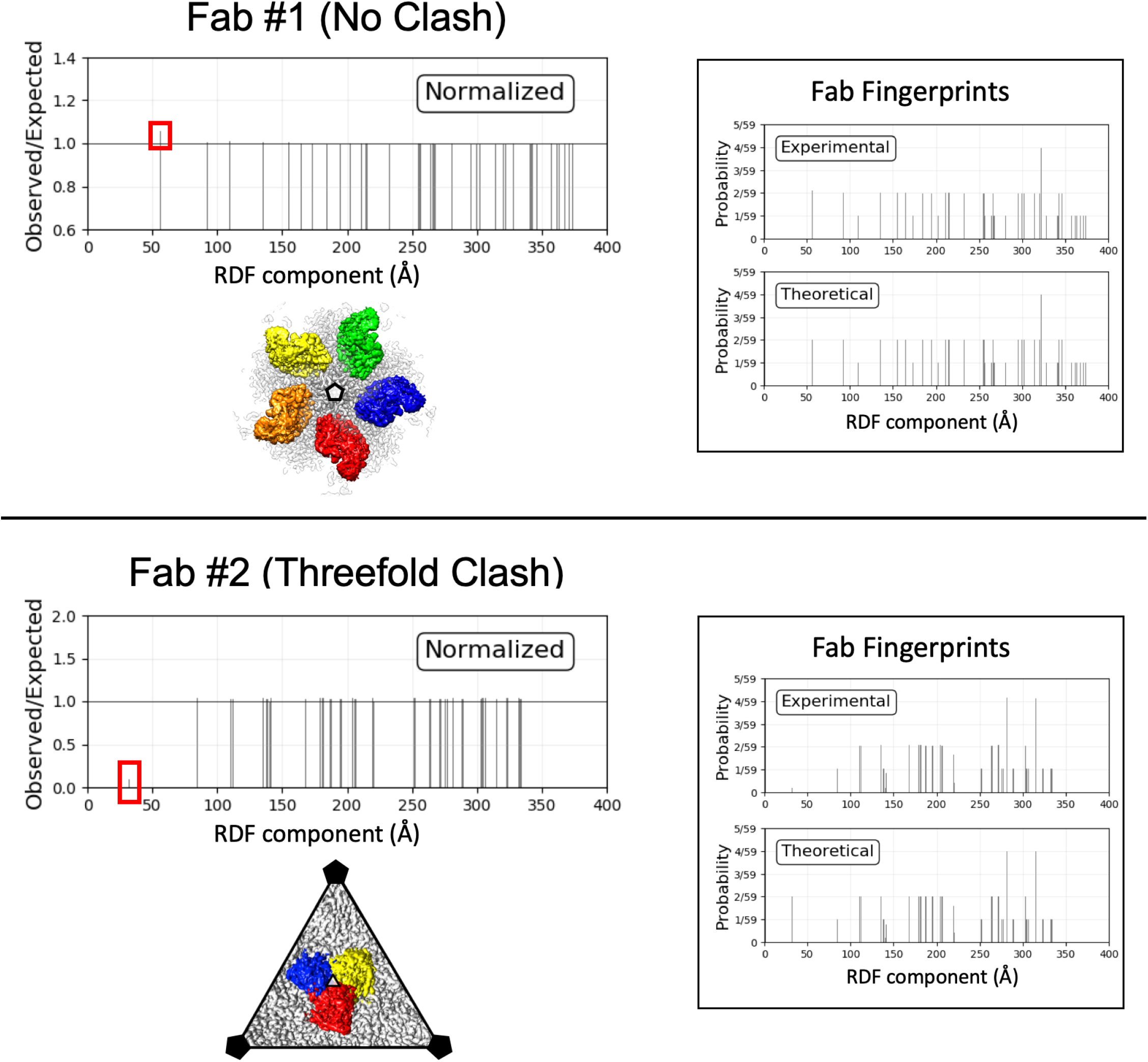
Further validation of RDFs for clash analysis. Two undersaturated enterovirus-Fab complexes are shown to demonstrate the impact of Fab binding geometry on cooperativity via RDF analysis. Fab #1 features no symmetry clash, with the closest interaction being the fivefold-relationship (top panel, Fab shown in red, orange, yellow, green, blue). Fab #2 features strong symmetry clash across the threefold symmetry axis (bottom panel, Fab shown in red, blue, yellow). The behavior of Fab #1 largely matched CPV-Fab 14, with a slight excess of the most proximal RDF component. Consistent with steric clash, the behavior of Fab #2 matched that of CPV-FabE. Enterovirus-Fab structures and corresponding biological analysis will be published separately. They are shown as further validation of Fab fingerprints in assessing binding cooperativity.

**S.Movie 1, 2.**
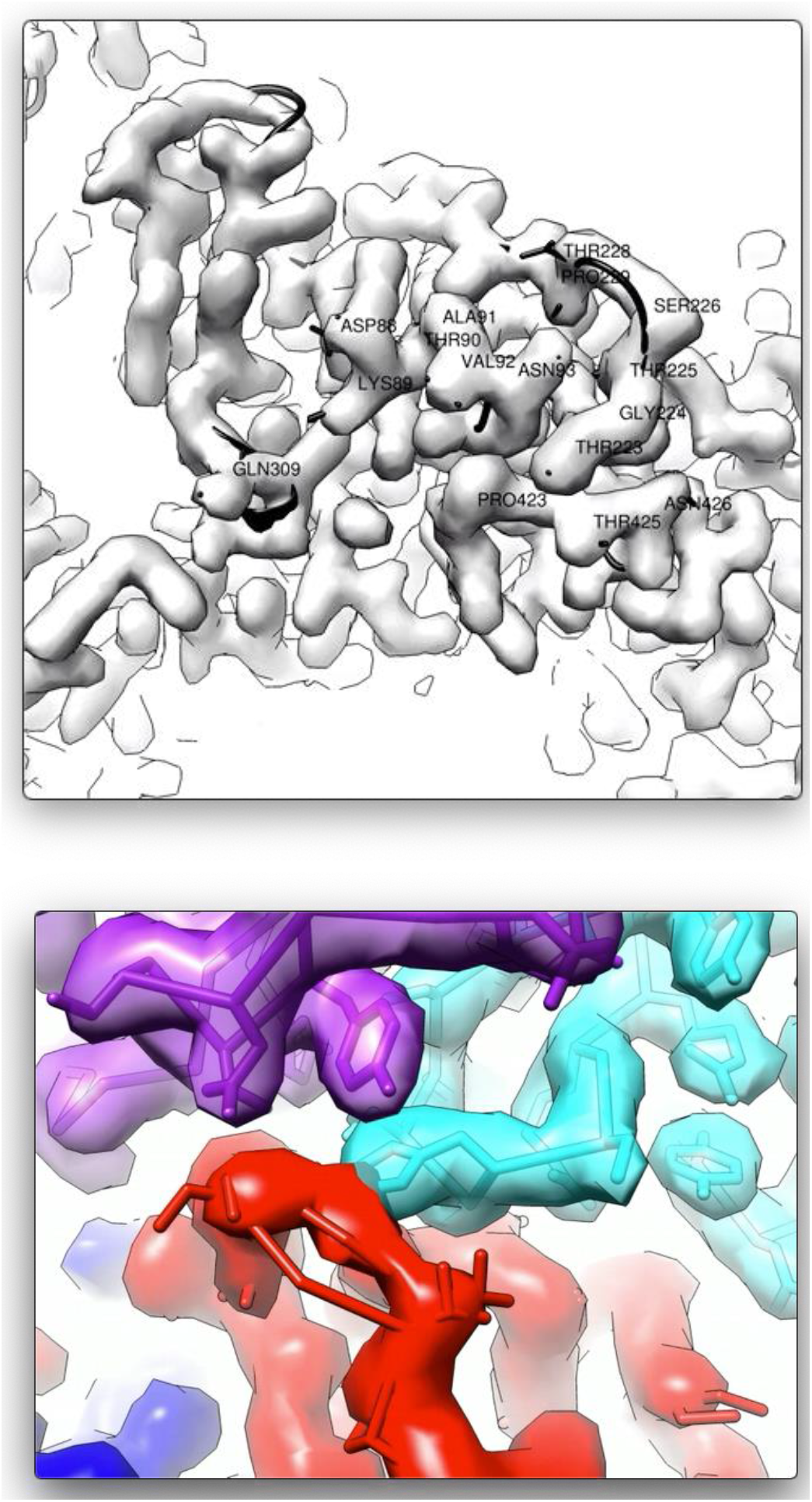
Occupied-Unoccupied epitope morph map. Morphing between the two particle-matched asymmetric maps suggests subtle hinging of the 228 loop on VP2. Key residues in the Fab 14 epitope are labeled. The Fab density was segmented away to provide this top-down view of the epitope. For side view, density is colored as in Fig. 2.

## References

1. Parrish CR, Kawaoka Y. 2005. The origins of new pandemic viruses: the acquisition of new host ranges by canine parvovirus and influenza A viruses. Annu Rev Microbiol 59:553–586.

2. Truyen U, Gruenberg A, Chang SF, Obermaier B, Veijalainen P, Parrish CR. 1995. Evolution of the feline-subgroup parvoviruses and the control of canine host range in vivo. J Virol 69:4702–4710.

3. Stucker KM, Pagan I, Cifuente JO, Kaelber JT, Lillie TD, Hafenstein S, Holmes EC, Parrish CR. 2012. The Role of Evolutionary Intermediates in the Host Adaptation of Canine Parvovirus. J Virol 86:1514–1521.

4. Hoelzer K, Parrish CR. 2010. The emergence of parvoviruses of carnivores. Vet Res 41:39.

5. Kaelber JT, Demogines A, Harbison CE, Allison AB, Goodman LB, Ortega AN, Sawyer SL, Parrish CR. 2012. Evolutionary Reconstructions of the Transferrin Receptor of Caniforms Supports Canine Parvovirus Being a Re-emerged and Not a Novel Pathogen in Dogs. PLoS Pathog 8:e1002666.

6. Allison AB, Organtini LJ, Zhang S, Hafenstein SL, Holmes EC, Parrish CR. 2016. Single Mutations in the VP2 300 Loop Region of the Three-Fold Spike of the Carnivore Parvovirus Capsid Can Determine Host Range. J Virol 90:753–767.

7. VanBlargan LA, Goo L, Pierson TC. 2016. Deconstructing the Antiviral Neutralizing-Antibody Response: Implications for Vaccine Development and Immunity. Microbiol Mol Biol Rev MMBR 80:989–1010.

8. Nelson CDS, Palermo LM, Hafenstein SL, Parrish CR. 2007. Different mechanisms of antibody-mediated neutralization of parvoviruses revealed using the Fab fragments of monoclonal antibodies. Virology 361:283–293.

9. Mietzsch M, Pénzes JJ, Agbandje-McKenna M. 2019. Twenty-Five Years of Structural Parvovirology. Viruses 11.

10. Tsao J, Chapman MS, Agbandje M, Keller W, Smith K, Wu H, Luo M, Smith TJ, Rossmann MG, Compans RW. 1991. The three-dimensional structure of canine parvovirus and its functional implications. Science 251:1456–1464.

11. Parrish CR, Carmichael LE. 1983. Antigenic structure and variation of canine parvovirus type-2, feline panleukopenia virus, and mink enteritis virus. Virology 129:401–414.

12. Strassheim ML, Gruenberg A, Veijalainen P, Sgro JY, Parrish CR. 1994. Two dominant neutralizing antigenic determinants of canine parvovirus are found on the threefold spike of the virus capsid. Virology 198:175–184.

13. Hafenstein S, Bowman VD, Sun T, Nelson CDS, Palermo LM, Chipman PR, Battisti AJ, Parrish CR, Rossmann MG. 2009. Structural comparison of different antibodies interacting with parvovirus capsids. J Virol 83:5556–5566.

14. Govindasamy L, Hueffer K, Parrish CR, Agbandje-McKenna M. 2003. Structures of host range-controlling regions of the capsids of canine and feline parvoviruses and mutants. J Virol 77:12211–12221.

15. Hueffer K, Govindasamy L, Agbandje-McKenna M, Parrish CR. 2003. Combinations of two capsid regions controlling canine host range determine canine transferrin receptor binding by canine and feline parvoviruses. J Virol 77:10099–10105.

16. Chang SF, Sgro JY, Parrish CR. 1992. Multiple amino acids in the capsid structure of canine parvovirus coordinately determine the canine host range and specific antigenic and hemagglutination properties. J Virol 66:6858–6867.

17. Wu H, Rossmann MG. 1993. The canine parvovirus empty capsid structure. J Mol Biol 233:231–244.

18. Simpson AA, Chandrasekar V, Hébert B, Sullivan GM, Rossmann MG, Parrish CR. 2000. Host range and variability of calcium binding by surface loops in the capsids of canine and feline parvoviruses. J Mol Biol 300:597–610.

19. Hafenstein S, Bowman VD, Sun T, Nelson CDS, Palermo LM, Chipman PR, Battisti AJ, Parrish CR, Rossmann MG. 2009. Structural comparison of different antibodies interacting with parvovirus capsids. J Virol 83:5556–5566.

20. Dunbar CA, Callaway HM, Parrish CR, Jarrold MF. 2018. Probing Antibody Binding to Canine Parvovirus with Charge Detection Mass Spectrometry. J Am Chem Soc 140:15701–15711.

21. Lindner JM, Cornacchione V, Sathe A, Be C, Srinivas H, Riquet E, Leber X-C, Hein A, Wrobel MB, Scharenberg M, Pietzonka T, Wiesmann C, Abend J, Traggiai E. 2019. Human Memory B Cells Harbor Diverse Cross-Neutralizing Antibodies against BK and JC Polyomaviruses. Immunity 50:668–676.e5.

22. Gurda BL, DiMattia MA, Miller EB, Bennett A, McKenna R, Weichert WS, Nelson CD, Chen W, Muzyczka N, Olson NH, Sinkovits RS, Chiorini JA, Zolotutkhin S, Kozyreva OG, Samulski RJ, Baker TS, Parrish CR, Agbandje-McKenna M. 2013. Capsid antibodies to different adeno-associated virus serotypes bind common regions. J Virol 87:9111–9124.

23. Agbandje M, McKenna R, Rossmann MG, Strassheim ML, Parrish CR. 1993. Structure determination of feline panleukopenia virus empty particles. Proteins 16:155–171.

24. Palermo LM, Hafenstein SL, Parrish CR. 2006. Purified feline and canine transferrin receptors reveal complex interactions with the capsids of canine and feline parvoviruses that correspond to their host ranges. J Virol 80:8482–8492.

25. Zhang K. 2016. Gctf: Real-time CTF determination and correction. J Struct Biol 193:1–12.

26. Scheres SHW. 2012. RELION: implementation of a Bayesian approach to cryo-EM structure determination. J Struct Biol 180:519–530.

27. Punjani A, Rubinstein JL, Fleet DJ, Brubaker MA. 2017. cryoSPARC: algorithms for rapid unsupervised cryo-EM structure determination. Nat Methods 14:290–296.

28. Emsley P, Cowtan K. 2004. Coot: model-building tools for molecular graphics. Acta Crystallogr D Biol Crystallogr 60:2126–2132.

29. Adams PD, Afonine PV, Bunkóczi G, Chen VB, Davis IW, Echols N, Headd JJ, Hung L-W, Kapral GJ, Grosse-Kunstleve RW, McCoy AJ, Moriarty NW, Oeffner R, Read RJ, Richardson DC, Richardson JS, Terwilliger TC, Zwart PH. 2010. PHENIX: a comprehensive Python-based system for macromolecular structure solution. Acta Crystallogr D Biol Crystallogr 66:213–221.

30. Chen VB, Arendall WB, Headd JJ, Keedy DA, Immormino RM, Kapral GJ, Murray LW, Richardson JS, Richardson DC. 2010. MolProbity: all-atom structure validation for macromolecular crystallography. Acta Crystallogr D Biol Crystallogr 66:12–21.

31. Ilca SL, Kotecha A, Sun X, Poranen MM, Stuart DI, Huiskonen JT. 2015. Localized reconstruction of subunits from electron cryomicroscopy images of macromolecular complexes. Nat Commun 6:8843.

32. Lauver MD, Goetschius DJ, Netherby-Winslow CS, Ayers KN, Jin G, Haas DG, Frost EL, Cho SH, Bator CM, Bywaters SM, Christensen ND, Hafenstein SL, Lukacher AE. 2020. Antibody escape by polyomavirus capsid mutation facilitates neurovirulence. eLife 9.

33. Sanchez-Garcia R, Gomez-Blanco J, Cuervo A, Carazo JM, Sorzano COS, Vargas J. 2020. DeepEMhancer: a deep learning solution for cryo-EM volume post-processing. bioRxiv 2020.06.12.148296.

34. Ludtke SJ, Baldwin PR, Chiu W. 1999. EMAN: semiautomated software for high-resolution single-particle reconstructions. J Struct Biol 128:82–97.

35. Callaway HM, Welsch K, Weichert W, Allison AB, Hafenstein SL, Huang K, Iketani S, Parrish CR. 2018. Complex and Dynamic Interactions between Parvovirus Capsids, Transferrin Receptors, and Antibodies Control Cell Infection and Host Range. J Virol 92.

36. Pettersen EF, Goddard TD, Huang CC, Couch GS, Greenblatt DM, Meng EC, Ferrin TE. 2004. UCSF Chimera--a visualization system for exploratory research and analysis. J Comput Chem 25:1605–1612.

37. Govindasamy L, Hueffer K, Parrish CR, Agbandje-McKenna M. 2003. Structures of host range-controlling regions of the capsids of canine and feline parvoviruses and mutants. J Virol 77:12211–12221.

38. Organtini LJ, Lee H, Iketani S, Huang K, Ashley RE, Makhov AM, Conway JF, Parrish CR, Hafenstein S. 2016. Near-Atomic Resolution Structure of a Highly Neutralizing Fab Bound to Canine Parvovirus. J Virol 90:9733–9742.

39. Chapman MS, Rossmann MG. 1995. Single-stranded DNA-protein interactions in canine parvovirus. Struct Lond Engl 1993 3:151–162.

40. Palermo LM, Hafenstein SL, Parrish CR. 2006. Purified feline and canine transferrin receptors reveal complex interactions with the capsids of canine and feline parvoviruses that correspond to their host ranges. J Virol 80:8482–8492.

41. Wang W, Ye W, Yu Q, Jiang C, Zhang J, Luo R, Chen H-F. 2013. Conformational selection and induced fit in specific antibody and antigen recognition: SPE7 as a case study. J Phys Chem B 117:4912–4923.

42. Sinha N, Smith-Gill SJ. 2005. Molecular dynamics simulation of a high-affinity antibody-protein complex: the binding site is a mosaic of locally flexible and preorganized rigid regions. Cell Biochem Biophys 43:253–273.

43. Rini JM, Schulze-Gahmen U, Wilson IA. 1992. Structural evidence for induced fit as a mechanism for antibody-antigen recognition. Science 255:959–965.

44. Parker JS, Parrish CR. 1997. Canine parvovirus host range is determined by the specific conformation of an additional region of the capsid. J Virol 71:9214–9222.

45. Llamas-Saiz AL, Agbandje-McKenna M, Parker JS, Wahid AT, Parrish CR, Rossmann MG. 1996. Structural analysis of a mutation in canine parvovirus which controls antigenicity and host range. Virology 225:65–71.

46. Xiao C, Rossmann MG. 2007. Interpretation of electron density with stereographic roadmap projections. J Struct Biol 158:182–187.

47. Rossmann MG, Palmenberg AC. 1988. Conservation of the putative receptor attachment site in picornaviruses. Virology 164:373–382.

